# Histone sequence variation in divergent eukaryotes facilitates diversity in chromatin packaging

**DOI:** 10.1101/2021.05.12.443918

**Authors:** Indu Patwal, Hien Trinh, Aaron Golden, Andrew Flaus

## Abstract

The histone proteins defining nucleosome structure are highly conserved in common model organisms and are frequently portrayed as uniform chromatin building blocks. We surveyed over 1700 complete eukaryotic genomes and confirm that almost all encode recognisable canonical core histones. Nevertheless, divergent eukaryotes show unrecognised diversity in histone sequences and offer an opportunity to observe the potential for nucleosome variation. Recombinant histones for *Plasmodium falciparum, Giardia lamblia, Encephalitozoon cuniculi* and *Leishmania major* were prepared alongside those for human, *Xenopus laevis* and *Saccharomyces cerevisiae*. All could be assembled into nucleosomes *in vitro* on sequences known to direct positioning with metazoan histones. *P. falciparum* histones refolded into very stable nucleosomes consistent with a highly regulated transcriptional programme. In contrast, *G. lamblia* and *E. cuniculi* histones formed less stable nucleosomes and were prone to aggregation as H3-H4 tetramers. Inspection of the histone fold dimer interface residues suggested a potential to form tetrasomal arrays consistent with polymerisation. DNA binding preferences observed using systematic evolution of ligands by exponential enrichment (SELEX) for human, *P. falciparum* and *E. cuniculi* histone octamers were highly similar and reflect a shared capability to package diverse genomic sequences. This demonstrates that nucleosomal organisation is retained across eukaryotes and can accommodate genome variation, but histone protein sequences vary more than commonly recognised to provide the potential for diversity of chromatin features.

**Significance statement:** It is widely assumed that eukaryotes package their genomes using equivalent nucleosome building blocks despite considerable variation in the composition and behaviour of cell nuclei. Our survey of available eukaryote genomes shows that histone proteins from divergent eukaryotes vary much more widely in sequence than is commonly recognised, even in histone fold dimer and DNA interaction interfaces. We demonstrate that divergent eukaryote histones nevertheless form nucleosomes on DNA sequences favoured in metazoans. These nucleosomes vary in stability but share broad DNA sequence preferences. This suggests that histone-dependent packaging does not constrain genome variation, and that chromatin behaviour can adapt by evolution of canonical core histone sequences in addition to other well-known mechanisms.

## Introduction

Canonical core histone proteins H2A, H2B, H3 and H4 direct the wrapping of genomic DNA into nucleosomes (1, 2). This structure is based on the specific and stable histone fold heterodimerisation of H2A-H2B and H3-H4 via ∼65 residue histone fold domains (HFDs) seeded by tight hydrophobic antiparallel packing of their long central α2 helices (3). Distinctive H3:H3 and H4:H2B 4 helix bundle (4HB) interfaces contributed by the HFD α2 and α3 helices then drive stacking into a spiralling octamer complex (4). High resolution structures have revealed that DNA wrapping around this octameric core involves over 100 protein-DNA contacts dominated by charge-charge interactions of basic histone sidechains and hydrogen bonding by the polypeptide backbone (5, 6). The many nucleosome crystal structures solved to date show very high structural homogeneity (7), albeit on a limited variety of DNA and histone protein sequences.

Overall, a well-defined arrangement of histone atoms must be scaffolded by the small HFD in order to direct the structure of the nucleosome and its DNA wrapping. The requirement for structural integrity has been interpreted as a strong selective pressure on the primary sequences of canonical histones, and it is often claimed that histones are amongst the most conserved proteins in eukaryotes. This is based on the very small numbers of residue changes between evolutionarily distant organisms such as animals, yeast and plants (8). Although there are only 2 amino acid differences between pea and human in the 62 residue H4 HFD (97% identity), there are in fact 12 differences between the most homologous pea and human H2B isoform HFDs (81% identity). Alanine scanning mutagenesis of *Saccharomyces cerevisiae* histones showed that mutation of only 40 of 228 (18%) non-alanine HFD residues are deleterious (9–11). It has also been demonstrated that reversion of only 5 of the 88 non-identical residues in the four human canonical core histones is required for robust growth in *S. cerevisiae* (12). This implies that at most 83% overall identity is required over the full length of the 4 histones, and none of the 5 reverted residues are in HFDs.

The major functions attributed directly to nucleosome packaging are compaction of the very long linear genomic DNA into microscopic three-dimensional nuclei, and negative regulation of cryptic transcription and self-replicating elements (13). Additional proposed roles for histones include regulatory participation, as epigenetic platforms, providing protection against mutagens, and imparting biophysical properties on cells via nuclear morphology.

Chromatin properties can be modulated locally and regionally by adjusting nucleosome composition using post-translational modifications (14) or replacement by histone variants (15), and global packaging can be altered by replacement of histones with other basic proteins such as in metazoan sperm (16) and dinoflagellates (17). Histone progenitors in some clades of Archaea exhibit the same overall DNA wrapping geometry as nucleosomes (18), although they form a continuous helical structures rather than discrete octameric units.

The phylogeny of eukaryotes is actively debated (19, 20) but it is generally accepted that a Last Eukaryotic Common Ancestor (LECA) existed some 1.9 billion years ago with a range of advanced features including a nucleus with histone-packaged chromatin (21). Diversification has led to a broad tree of eukaryotes with fungi and animals grouped together and plants branching earlier (20). In addition, there are a large number of evolutionarily more distant and diverse unicellular eukaryotes referred to as protists. These include ciliates, apicomplexans and amoeba which have been studied because of their biomedical implications as infective human parasites.

By surveying histones in sequenced genomes of over 1700 eukaryotes we observed that even the most distantly related eukaryotes encode canonical core histone sequences. The histones from these divergent eukaryotes have much less sequence identity to human histones than commonly studied model organisms such as *Xenopus laevis* or *S. cerevisiae* that are widely used in biochemical experiments. To investigate the effect of sequence variation on nucleosome structure and stability we prepared sets of recombinant histones for 7 different organisms and assembled them into nucleosomes. We then observed *in vitro* nucleosome stability, and DNA sequence preferences using systematic evolution of ligands by exponential enrichment (SELEX). Our findings are consistent with an unexpected variability in canonical core histone sequences that facilitates adaptability for both genome sequences and chromatin packaging.

## Results

### Assessment of histone diversity in eukaryotes

To quantitate the ubiquity and diversity of canonical core histones we surveyed 1725 predicted eukaryotic proteomes in the Ensembl Genomes resource comprising 305 vertebrates, 117 non-vertebrate metazoans, 96 plants, 1014 fungi and 237 protists. Custom BLAST searches of individual genomes using human reference HFD sequences followed by global alignment to these HFD revealed the proteomes of 99% of sequenced eukaryotes have been predicted to encode at least 1 canonical core histone. All four histones can be identified in 93% of genomes with at least 30% identity to human HFDs (supplementary figure 1). Allowing for incompleteness in sequencing, assembly, gene calling and the uneven phylogeny of whole genome sequencing, this provides a direct validation of the near ubiquity of the four core histones across eukaryotes.

**Figure 1:**
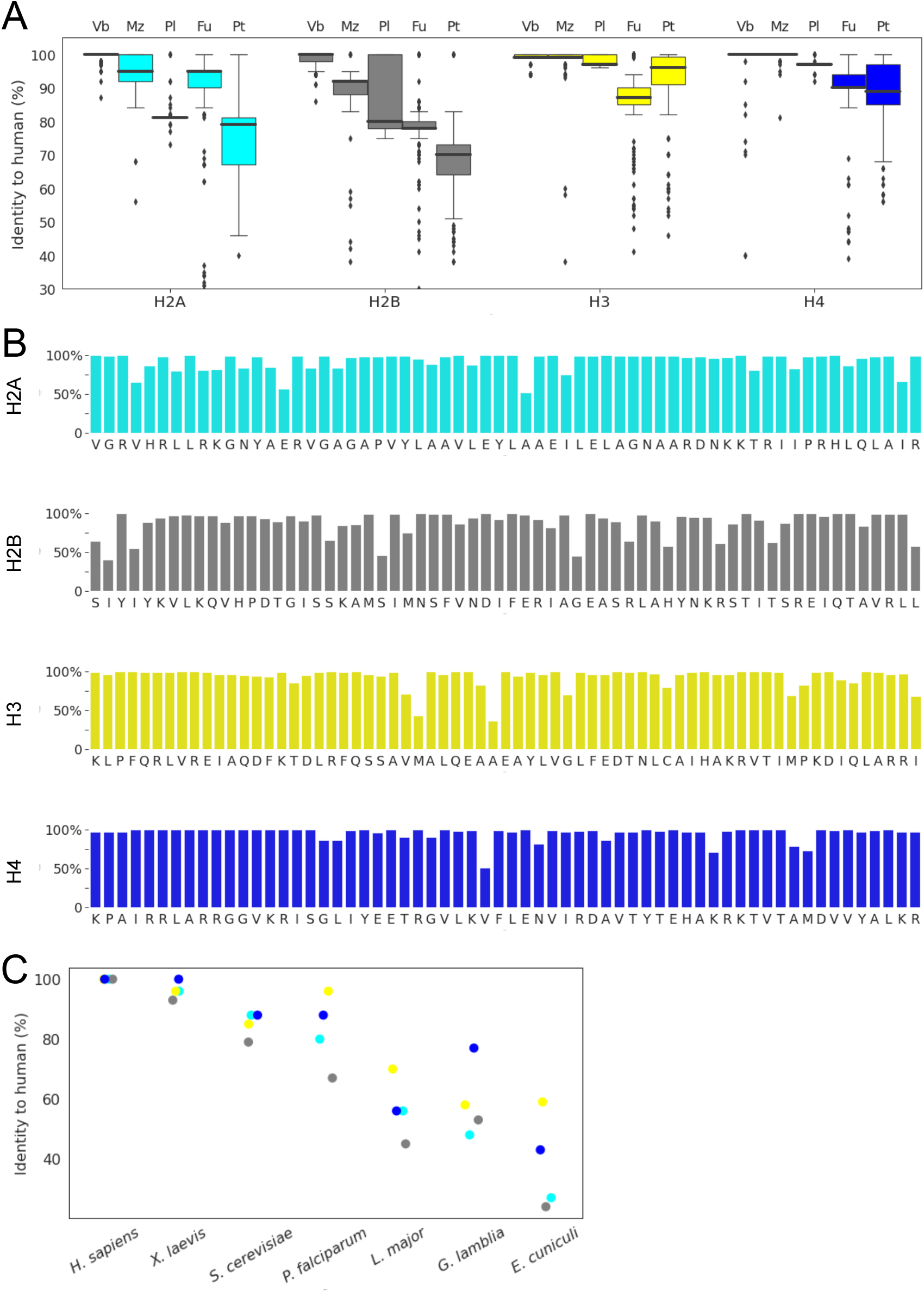
Phylogenetic diversity of core histones. A. Boxplot of highest identity to human reference H2A (cyan), H2B (grey), H3 (yellow) and H4 (blue) for 1208 unique species in phylogenetic groupings of vertebrates (Vb), non-vertebrate metazoans (Mz), plants (Pl), fungi (Fu) and protists (Pt) assigned by Ensembl Genomes. B. Conservation by residue across histone fold domains (HFDs) for 21,571 histones from 1769 genomes with ≥40% identity to human, indexed by human reference sequence with gaps removed. C. Identity of highest identity histones from selected organisms to human reference sequences as used for subsequent biochemical analysis.

We then calculated the sequence identity to human HFDs for the best matching sequence from the 1208 unique species in Ensembl Genomes, taking the closest match as representative for multiple subspecies. Plotting the best HFD identity for each unique species across the phylogenetic groupings reveals that histones are more diverse than often assumed, including a consistent proportion of lower homology outliers (figure 1A). The mean identities to the human reference HFDs across eukaryotes are 91%, 83%, 92% and 93% for H2A, H2B, H3 and H4 respectively. The distributions suggest that more diverged eukaryotes have more divergent histones relative to human. Within the vertebrate and plant phylogroups the core histones are almost identical in all species whereas fungi and protists have greater spreads in sequence diversity. Figure 1A also shows that HFDs of tetramer histones H3 and H4 are more highly conserved than dimer histones H2A and H2B. Additional broad trends include divergence of H2A and H2B HFDs in plants, and relative conservation of H2A compared to H2B HFDs in fungi.

We next plotted the frequencies of consensus residues at each position across the four respective HFDs for the 21571 total histones identified from all 1769 genomes with at least 40% identity to human HFDs (figure 1B). 62 of 255 positions (24%) across the four core histone HFDs are <90% conserved, reflecting tolerance for residue changes at most sites in the HFD structure. Patterns of variation can be observed in the fungal and protist sequences (supplementary figure 2) including conservation of α3 helices and of H3 and H4 L2 loops versus variability of H2B and H4 α2 helices and H2A and H2B L1 and L2 loops. The observed patterns of variation provide insights into motifs where nucleosome stability is modulated and suggests that variability in nucleosome structure and stability is being explored in eukaryote evolution.

**Figure 2:**
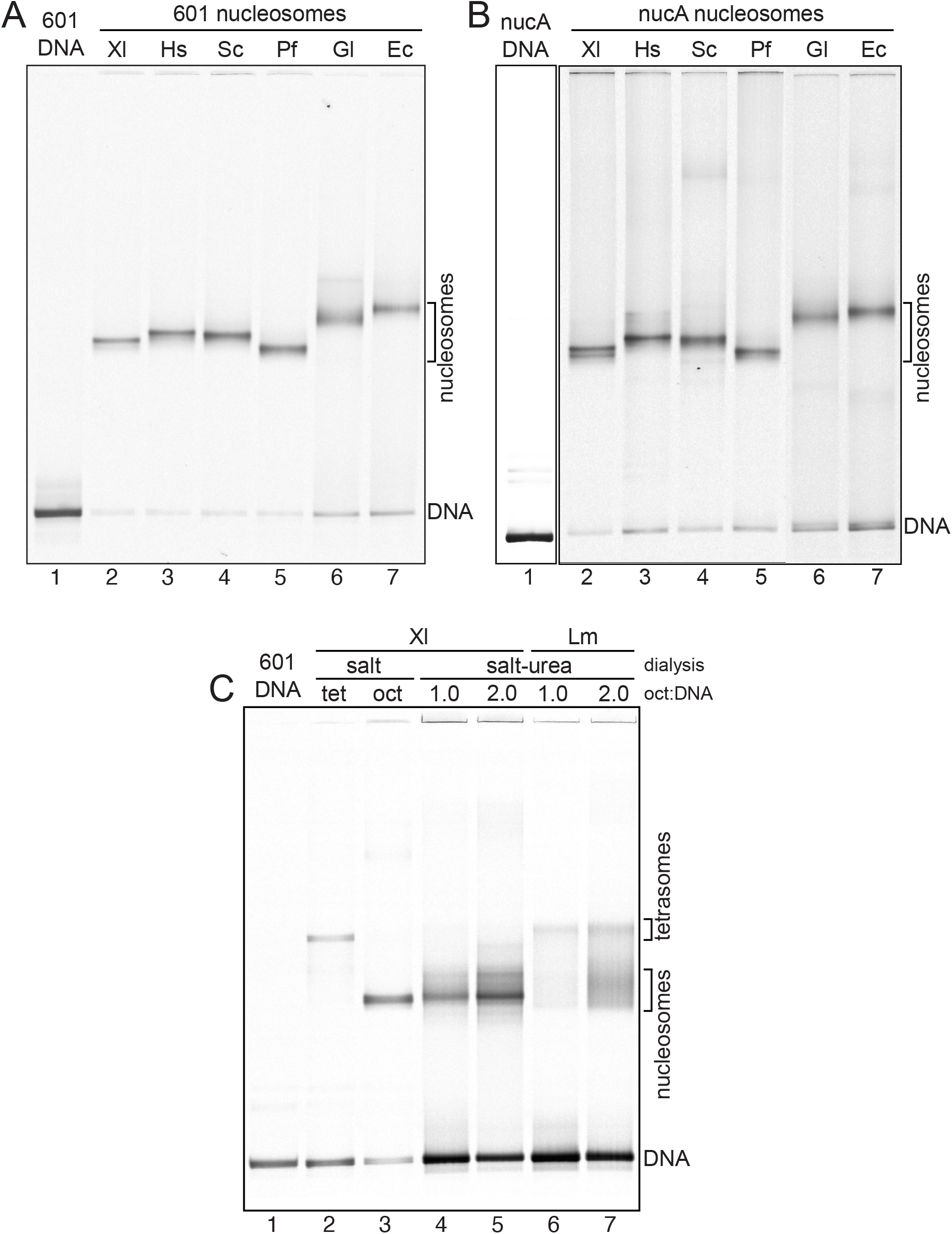
Nucleosomes assembled on positioning DNA fragments. A. Native PAGE of 147 bp 601 free DNA (lane 1) and nucleosomes assembled with *X. laevis* (Xl), *H. sapiens* (Hs), *S. cerevisiae* (Sc), *P. falciparum* (Pf), *G. lamblia* (Gl), and *E. cuniculi* (Ec) in lanes 2-7. B. Native PAGE with 147 bp MMTV nucA and salt dialysis assembly as in A. C. 147 bp 601 DNA free (lane 1) assembled with *X. laevis* (Xl) tetrasome (lane 2) and octamer (lane 3) by salt dialysis, and *X. laevis* and *L. major* (Lm) histones assembled at 1:1 and 2:1 octamer to DNA ratio by salt-urea dialysis in lanes 4-7.

### In vitro assembly of nucleosomes representing divergent eukaryotes

To test the implications of this sequence diversity on nucleosome structure and stability we identified the orthologous sequences with highest identity to human histones in the genomes of *Plasmodium falciparum, Giardia lamblia, Leishmania major* and *Encephalitozoon cuniculi*. The identities to human sequences for the HFDs of these histones range from 97% for *P. falciparum* H3 to 24% for *E. cuniculi* H2B (figure 1C and supplementary figure 3).

**Figure 3:**
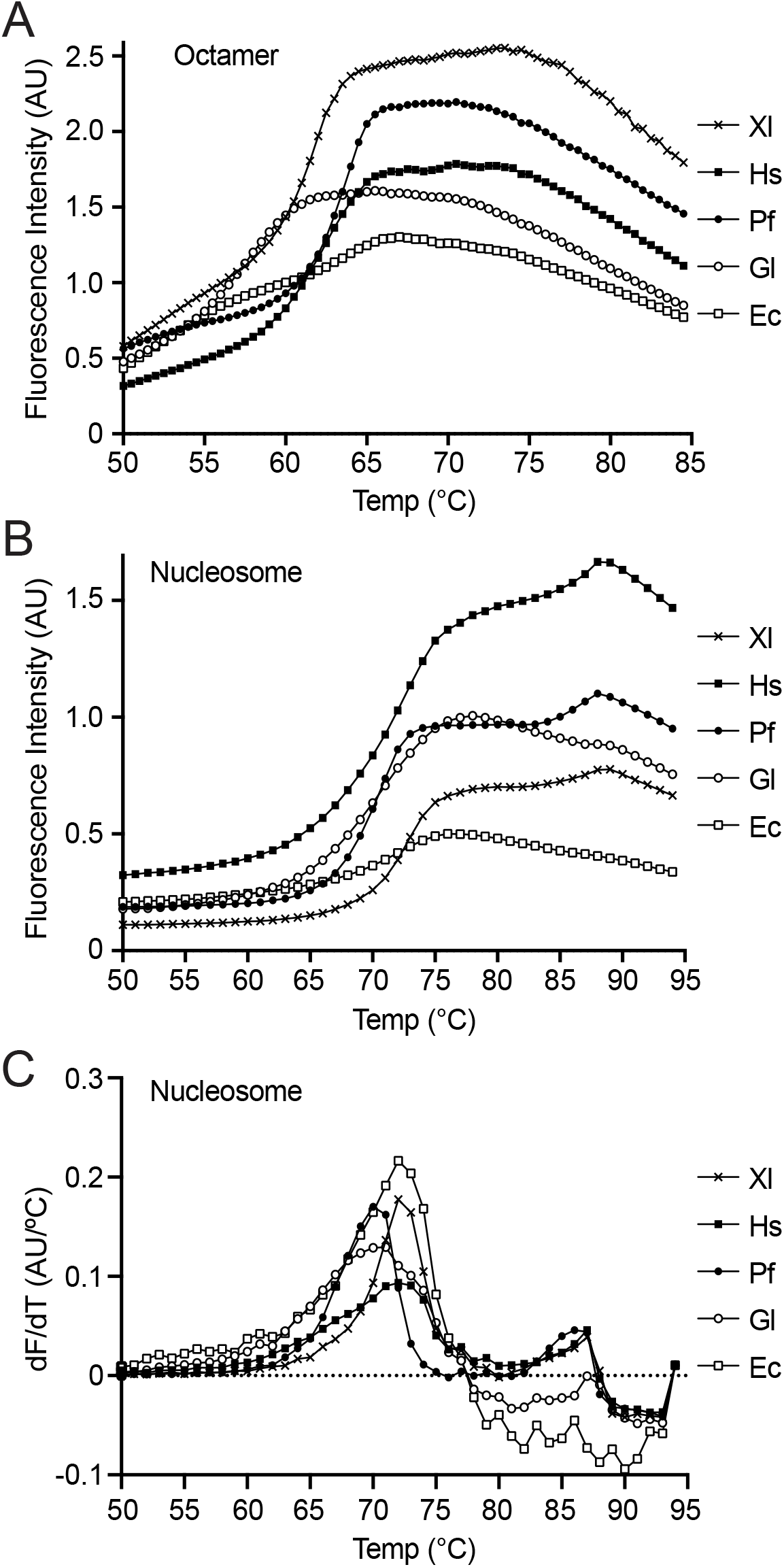
Octamer and nucleosome thermal stability. A. Thermal stability assay of *X. laevis* (Xl), *H. sapiens* (Hs), *S. cerevisiae* (Sc), *P. falciparum* (Pf), *G. lamblia* (Gl), and *E. cuniculi* (Ec) octamers. B. Thermal stability assay for nucleosomes assembled with octamers in A on 147 bp 601 DNA. C. Derivative plot for data in B.

These unicellular organisms were selected because information is available on their molecular biology, they have a range of genome properties, and they are agents of human disease. Compared to higher animals such as human with 3000 Mbp DNA and 41% GC, *P. falciparum* is an apicomplexan protozoan parasite with intracellular and extracellular phases that can cause malaria. It has a 23 Mbp genome that is only 19% GC. *G. lamblia*, also referred to as *G. intestinalis*, is a diplomonad protozoan extracellular parasite that can cause giardiasis. It has a 12 Mbp genome that is 49% GC. *L. major* is an intracellular euglenozoan protozoa parasite that can cause leishmaniasis. It has a 33 Mbp genome that is 60% GC. Finally, *E cuniculi* is an intracellular fungal parasite that can cause microsporidiosis. It has a minimal 3 Mbp genome that is 47% GC. For comparison, the budding yeast *S. cerevisiae* has a 12 Mbp genome that is 38% GC.

Genes encoding the selected histones for each organism were optimised and synthesised for recombinant expression in *Escherichia coli* then all 16 histones were expressed and purified in multi-milligram quantities by standard methods (supplementary figure 4). In addition, core histones for human, *Xenopus laevis* and *S. cerevisiae* were prepared for comparison because high resolution structures of these nucleosomes are available.

**Figure 4:**
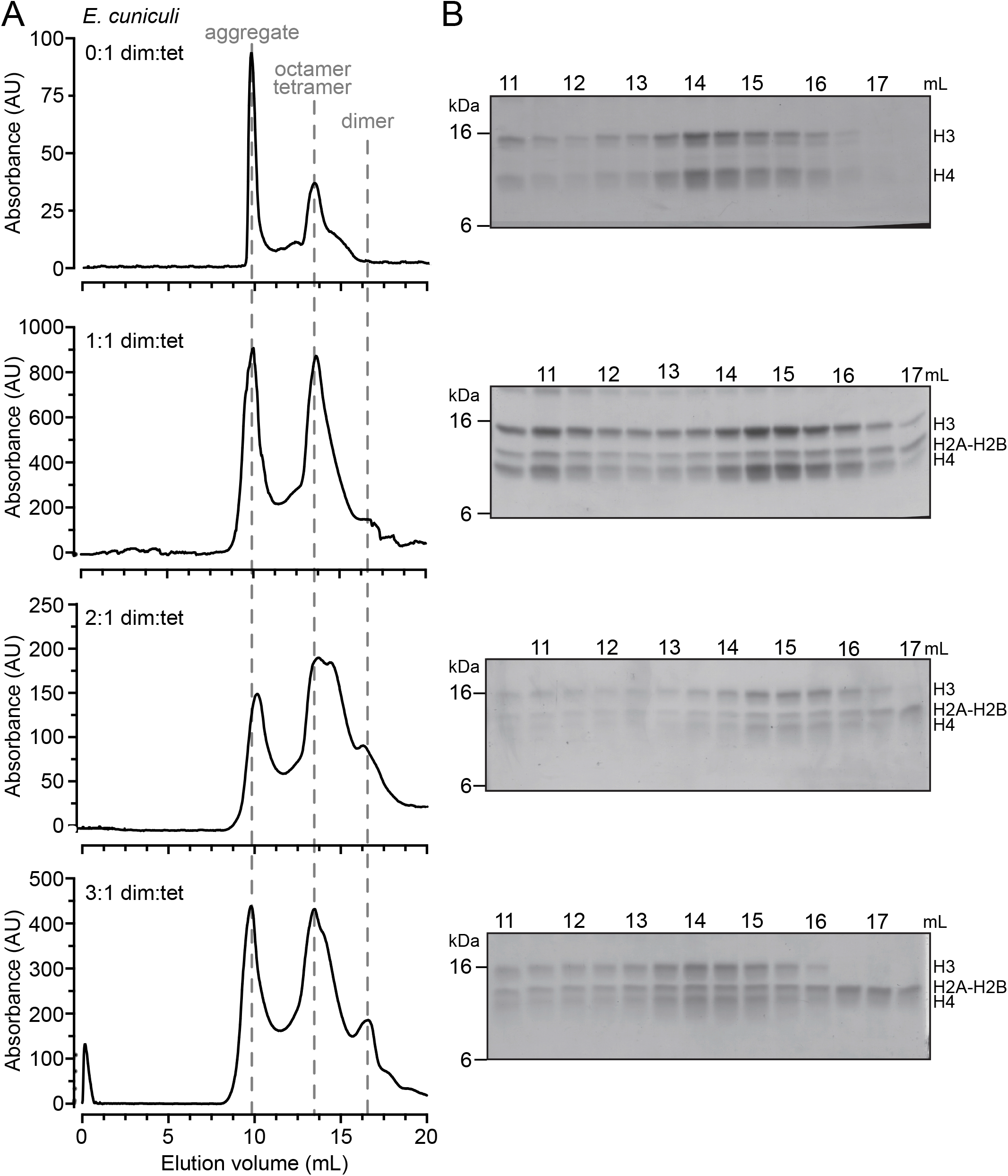
Titration of *E. cuniculi* tetramer with increasing dimer. A. Gel filtration chromatograms of *E. cuniculi* (H3-H4)_2_ tetramer with indicated molar ratios of H2A-H2B dimer after refolding in 2 M NaCl buffer. B. SDS PAGE of fractions from A stained with Coomassie.

Purified recombinant histones were unfolded using a strong chaotrope then dialysed into high ionic strength buffer to refold octamers which were purified by gel filtration chromatography. *P. falciparum* histones readily formed well resolved octamers equivalent to human and *X. laevis* complexes (supplementary figure 5). In contrast, *G. lamblia* and *E. cuniculi* histones consistently gave low octamer yields because a large proportion of histones aggregated during refolding despite all optimisation efforts. *L. major* histones could not be refolded into octamers by standard methods in our hands.

Purified octamers were then mixed with 147 bp DNA containing either the mouse mammary tumour virus nucleosome A (nucA) or Widom 601 (601) positioning sequences and assembled into nucleosomes by stepwise salt gradient dialysis. *L. major* nucleosomes could be assembled with lower efficiency by an alternative method involving mixing of individual histones and DNA in urea at high ionic strength then stepwise dialysis to low salt. The resulting nucleosome core particles were analysed by high resolution native PAGE (figure 2).

Nucleosome assembly in parallel for the 6 octamers reproducibly displayed the same relative native PAGE electrophoretic mobilities on both 147 bp 601 and nucA DNAs (figure 2). *P. falciparum* histones assembled into nucleosomes that migrate slightly more rapidly than the widely used *X. laevis* histones, whereas the *G. lamblia* and *E. cuniculi* histones formed significantly slower migrating particles. Nucleosomal bands extracted from native PAGE gels contained tryptic peptides representing all 4 core histones (data not shown). We then performed solution Fe/EDTA footprinting on all 6 histone octamers assembled with 601 DNA to confirm the appropriate wrapping of DNA around the nucleosomes and observed effectively identical DNA protection on both strands at all superhelical locations indicating equivalent periodicity (supplementary figure 6).

### Divergent nucleosomes vary in stability

The differences in native PAGE mobility do not correlate with differences in mass or net charge (supplementary table 2), so migration suggests that nucleosomes have differential hydrodynamic radii as a consequence of histone sequence variation. Structural studies of *S. cerevisiae* nucleosomes showing destabilised interactions between H2A L1 loops due to changes in hydrogen bonding of H2A N38 and E41 (22) are consistent with increased internal dynamics resulting in an enlarged mean hydrodynamic radius and slower migration. The L1 loop is the most variable motif within the HFD (supplementary figure 2), and human nucleosome behaviour could also be influenced by the adjacent H2A A40S change from *X. laevis. P. falciparum* contains a novel positively charge pairing of H2A K38 K41 reminiscent of L1 loop stabilisation in macroH2A (23) that could lead to reduced internal dynamics and faster native PAGE migration.

In contrast, the *E. cuniculi* and *G. lamblia* nucleosomes migrated considerably slower in native PAGE suggesting a major differentiation of chromatin properties. To compare their stability directly we assembled nucleosomes on 601 DNA and performed dye binding thermal stability assays (24). Under our assay conditions, *P. falciparum* and *G. lamblia* nucleosomes release H2A-H2B dimers approximately 2ºC earlier than *X. laevis*, human and *E. cuniculi* at 72ºC consistent with increased dynamics (figure 3A). In addition, *E. cuniculi* and *G. lamblia* exhibit a very unusual profile for tetrasome release over a broad range from 75ºC instead of the expected sharp unfolding above 85ºC (24). This led us to investigate the stabilisation of the *E. cuniculi* tetramer.

### Octamer formation differs for histones of divergent organisms

Octamer refolding of *E. cuniculi* and *G. lamblia* histones reproducibly resulted in significant loss of material as insoluble aggregates. Gel filtration chromatography of remaining soluble material revealed that aggregates are dominated by H3 and H4, leaving excess free soluble H2A-H2B dimers (data not shown). To investigate this further we assembled *E. cuniculi* H3 and H4 into tetramers at a range of salt concentrations from 0.5 to 2.0 M NaCl with *X. laevis* histones as a control. *X. laevis* H3-H4 assembly resulted in aggregation of H3 at low salt but efficient tetramer production at higher salt concentrations with limited aggregation, whereas high levels of precipitation of both H3 and H4 occurred for *E. cuniculi* especially at increased ionic strength (supplementary figure 7). This is consistent with *E. cuniculi* H3-H4 tetramers associating into oligomeric and insoluble polymeric complexes driven by hydrophobic interactions that are enhanced under high ionic strength conditions.

Titration of increasing molar ratios of H2A-H2B dimer into the *E. cuniculi* H3 and H4 assembly reactions led to significant recovery of soluble octamer complexes (figure 4). This rescue can be rationalised by the H2A-H2B dimer blocking an interface responsible for aggregation. Comparison of the 4HB interfaces of *X. laevis* and *E. cuniculi* shows a change in the chemical nature of H4 4HBs by substitution of hydrophobic functional groups leading to a more hydrophobic nature for *E. cuniculi* H4 residues whereas the H3 interface residues are largely unchanged (supplementary figure 8). We speculate that this H4 variation creates a capability to form extended H3-H4 polymers (supplementary figure 8C) reminiscent of the equivalent archaeal chromatin structure which forms a continuous superhelix by packing of the 4HB interface (18) yet is inherently dynamic (25).

To investigate the potential significance of tetramer polymerisation we attempted to assemble the limited purified amounts of *E. cuniculi* tetramers on MMTV-derived DNA fragments of lengths from 147-252 bp. These were unable to efficiently form complexes with more than one tetramer whereas control titrations of *X. laevis* tetramers on the same DNA resulted complexes with migration in native gels of 1-3 tetramers as expected (supplementary figure 9A). Inspection of the MMTV DNA sequences revealed regional variation in GC content so we constructed fragments spanning the MMTV nucA and nucB region and observed that *E. cuniculi* tetramers appeared to assemble more efficiently on the fragment with lowest GC content (supplementary figure 9B and 9C) pointing to an influence of DNA sequence on assembly of these chromatin complexes.

### SELEX reveals strong nucleosome formation for 601 and two novel sequences

We performed SELEX on a human mononucleosomal DNA library using 3 different octamers to investigate the sequence preferences of the divergent histones. A library of ∼191 bp fragments was constructed by ligating a 22 bp adapter onto both ends of 147 bp mononucleosomal fragments obtained from human HEK293 chromatin after extensive micrococcal nuclease digestion. The input library for SELEX had 4 × 10^5^ unique sequences, limited by tight mononucleosome selection and suboptimal adapter ligation efficiency. Nucleosomes were assembled in parallel using human, *P. falciparum* and *E. cuniculi* histone octamers at a molar ratio of 1:10 to the DNA library. The assembled complexes were then separated by native PAGE and nucleosomal DNA bands were extracted and reamplified, and this process was repeated for 6 cycles of SELEX. The outputs of rounds 1, 3 and 6 were submitted for next generation sequencing and selected DNAs were identified using a pipeline involving quality control of reads, mapping of paired reads to the human genome, and filtering for length and mapping quality.

Enriched sequences were identified by monitoring their excess over power law expectation (26). The enrichment was complete by SELEX cycle 6 for both human and *P. falciparum* octamers resulting in the identification of 9 and 5 unique sequences respectively (table 1, supplementary table 3) and enrichment was near completion for *E. cuniculi* octamers after 6 cycles with 15 sequences identified. The selected sequences were overlapping for all octamers, and were dominated by 601, a novel *ANK1*-*POU6F2* fusion, and a fragment from *INHBB* exon 2.

**Table 1:** Mapped DNA fragment counts from nucleosome SELEX. Total properly mapped paired end read counts by Bowtie2 and samtools after quality control, counts per million total fragments (cpM) to 147 bp regions for 601, MMTV nucA, *ANK1-POU6F2* fusion and for human genome (GRCh37), and within human genome fragments for *INHBB* exon 2 and *PSD2* intron 4 fragments (S1 and S2 respectively in supplementary tables 3-4).

| Octamer in nucleosome | SELEX round | Total reads after QC | Mapping to 601 (cpM) | Mapping to nucA (cpM) | Mapping to *ANK1-POU6F2* fusion (cpM) | Mapping to human genome (cpM) | Mapping in human genome to *INHBB* exon 2 (S1, cpM) | Mapping in human genome to *PSD2* intron 4 (S2, cpM) |
| --- | --- | --- | --- | --- | --- | --- | --- | --- |
| - | pre-Library | 25,336,620 | 37 | 86 | 6 | 933,287 | 13 | 7 |
| - | Library | 1,207,124 | 216,102 | 126,854 | 32 | 616,373 | 530 | 69 |
| Human | 1 | 12,713,025 | 18,448 | 1,592 | 8,572 | 934,538 | 28,211 | 18,277 |
| Human | 3 | 14,493,439 | 430,126 | 59 | 129,545 | 536,645 | 758,476 | 84,700 |
| Human | 6 | 15,671,881 | 706,865 | 0 | 36,346 | 259,073 | 904,798 | 93,072 |
| *P. falciparum* | 1 | 12,064,019 | 16,981 | 800 | 1,300 | 944,484 | 22,842 | 5,216 |
| *P. falciparum* | 3 | 16,369,858 | 601,156 | 44 | 23,619 | 373,600 | 909,399 | 11,505 |
| *P. falciparum* | 6 | 12,947,118 | 294,642 | 0 | 1,436 | 674,929 | 999,613 | 173 |
| *E. cuniculi* | 1 | 11,882,054 | 2,545 | 2,319 | 10,474 | 908,418 | 35,916 | 7,820 |
| *E. cuniculi* | 3 | 13,556,692 | 124,866 | 111 | 184,664 | 825,663 | 678,815 | 83,340 |
| *E. cuniculi* | 6 | 14,085,343 | 282,079 | 0 | 224,747 | 670,009 | 767,233 | 229,581 |

Adventitious introduction of 601 and nucA into the initial library provided an internal control. 601 DNA was rapidly selected and comprised 12-60% of all fragments within 3 cycles (table 1), consistent with the original isolation of the 601 sequence by SELEX (27). Conversely, MMTV nucA that entered the library with similar abundance to 601 was represented by less than 1% of sequences after 3 cycles and lost at cycle 6.

A novel *ANK1*-*POU6F2* fusion fragment was enriched to levels similar to 601 after human and *E. cuniculi* octamer SELEX cycle 3 although not as highly for *P. falciparum*, and it remained with *E. cuniculi* even after 6 cycles (table 1). This fragment comprised a fusion of intron sequences from the human *ANK1* and *POU6F2* genes with a 4 bp overlap at their junction. The continuous sequence does not exist in the human reference genome or a recently published high depth HEK293 assembly (28), and reads crossing the fusion junction are not found in the HEK293 genome sequencing depositions. However, reads across the fusion boundary do exist in the progenitor HEK293 micrococcal nuclease (MNase) digested pre-Library input DNA prior to adapter ligation in our SELEX library construction. This suggests the high affinity *ANK1*-*POU6F2* nucleosome binding sequence arose from genome rearrangements in the lineage of HEK293 cells before MNase, and this cell line is known to have a heterogeneous and unstable karyotype (29). The *ANK1*-*POU6F2* fragment can be assembled into mononucleosomes equivalent to 601 and nucA, with thermal unfolding similar to nucA but less stable than 601 (supplementary figure 10).

The most strongly enriched sequence mapping directly to the human genome arises from a site at the 3’ end of the coding region of the *INHBB* growth factor (table 1). *INHBB* is a Polycomb target (30) and the identified site is adjacent to an ATAC-seq accessible peak that binds the Runx2 pioneer transcription factor in mouse (31). A sequence from intron 4 of the *PSD2* gene was also notably enriched only in *E. cuniculi*, where it was represented at a similar order of magnitude to 601, *ANK1-POU6F2* and *INHBB*.

The similarities and differences in selection appear to represent a common foundation of *in vitro* assembly but with moderate differentials in affinities between the three species octamers. The 601 sequence was originally isolated using chicken octamer so it is unsurprising that this was dominant in human, whereas the *INHBB*-derived fragment was moderately more abundant for *P. falciparum* and *E. cuniculi* octamers (table 1). *E. cuniculi* octamers selection was slower to converge in SELEX and showed qualitatively similar preference for the *ANK1-POU6F2* fusion and *PSD2*-derived sequence as well as 601 and *INHBB*. Overall this reflects broadly similar preferences of the highly diverged histone octamers leading to selection of a small number of apparently unrelated DNA sequences, but with some variation between them under the experimental conditions. Any biological relevance for the specific sequences selected is unknown since the 601 sequence does not exhibit dominant properties when incorporated into genomes (32).

Despite this common selection, the sequences have no obvious homology and analysis of nucleotide abundance in sequences enriched by SELEX did not reveal any remarkable features (supplementary table 4). Almost all selected sequences have G/C compositions in the range 50-60%, which is moderately above the mean for the genomes packaged by the respective histones and for the input HEK293-derived mononucleosomal DNA library. However, the nucleotide compositions of the SELEX-selected sequences are seldom in the lowest or highest decile for the frequency of G/C (supplementary table 4), any mono or dinucleotide combination of the respective genome or the input library (data not shown). Likewise, analysis of dinucleotide periodicities did not reveal any common repetitive dinucleotide pattern (supplementary table 4).

## Discussion

Detailed bioinformatic investigations have identified histone fold containing proteins in all three domains of life as well as viruses, and catalogued considerable amplification of copy number and diversification of function for this motif (33, 34). This has led to a common but untested claim that canonical core histones are nearly universal in eukaryotes, and an assumption that their ubiquity and fundamental role in genome function implies their sequences are highly conserved.

We have confirmed that homologues with at least 40% identity in the HFD to each of the four human canonical core histones are encoded in approximately 90% of all currently sequenced eukaryotic genomes, and at least one of the histones can be detected above this threshold in 99% of genomes. Although genome sequencing of eukaryote diversity is highly uneven and automated gene calling is imperfect, this strongly supports the assumption of near ubiquity of histones without precluding isolated exceptions such as dinoflagellates (17). Nevertheless, we also observed that the core histone HFD sequences are more divergent than often assumed with mean identity between human and the best HFD matches for the 237 protists of only 67-91%.

To test the effect of this sequence variation we expressed and attempted to assemble both histone octamers and nucleosomes from a number of species with HFD identities of as low as 24-44% for the most conserved homologues in *E. cuniculi*. Histones from all of these species with the exception of *L. major* could form histone octamers as judged by gel filtration chromatography, and all could form mononucleosomes as judged by native PAGE and hydroxy radical footprinting including for nucleosomes composed of histones from the highly divergent *E. cuniculi*. This provides experimental support for the textbook model of nucleosomes as the universal building block of eukaryotic chromatin.

The ability of nucleosome structure to tolerate histone sequence variation at lesser levels has been observed in functional screening of point mutations and species-specific gene substitutions (9–12), and is consistent with general principles of protein folding and stability. The ubiquity of 4 core histones suggests that LECA likely used nucleosomes for genome packaging. However, this does not necessarily imply that LECA had also evolved all the functional dependencies seen in experimental model organisms. The conservation within phylogenetic groups such as animals or plants is likely to be driven by the role of nucleosomes as the substrate and scaffold for specific genomic mechanisms such as substrates for epigenetic marks or remodelling. The consequences of such additional requirements are reflected in features such as the oncohistones and specialised histone variants.

Our results support a hypothesis that the ability to facilitate diversification of biochemical features of chromatin is an advantage of nucleosomal packaging. Firstly, histones octamers from the range of species we tested accommodated stable binding of the artificial 601 and mouse-derived nucA that do not exist in the genomes they package. In fact, MMTV nucA spans a fusion of retroviral and native mouse sequences (35), and the *ANK1-POU6F2* fusion is probably a novelty of genome instability. In SELEX, *P. falciparum* and *E. cuniculi* octamers enriched strongly for the same 601, *ANK1-POU6F2* and *INHBB* sequences preferred by human octamers. None of the selected sequences were extreme in nucleotide composition or periodicity, although they did show elevated GC% consistent with models for sequence preference (36, 37). Interestingly, *P. falciparum* octamers appeared to show somewhat tighter selectivity and *E. cuniculi* showed looser sequence constraints judged by SELEX enrichment. Together, this is consistent with nucleosomes having the capability to package effectively all genomic sequences in order to avoid restricting genetic diversity, but still allowing the potential for phylogenetic groups to evolve individual constraints on sequences to be packaged such as exclusion of A/T-rich regions (38) or DNA methylation (39).

Secondly, the histone octamers from the species we tested exhibited a range of stabilities when assembled into nucleosomes and assayed by native PAGE or thermal denaturation. In our hands nucleosomes containing *P. falciparum* octamers were more compact although they were not more thermally stable than with human octamers, consistent with previous observations of these nucleosomes (40) and offering a potential explanation for the tightly regulated genetic program in this organism (41, 42) by rigid yet malleable nucleosomes. One possible contribution to rigidity is through variation in H2A L1-L1 interactions which involve one of the most variable motifs within the HFD (supplementary figure 2). In contrast, *E. cuniculi* as well as *G. lamblia* nucleosomes were significantly less compact in native PAGE.

Finally, both *E. cuniculi* and *G. lamblia* histone octamers are prone to aggregation, particularly of H3-H4 tetramers. We demonstrated that *E. cuniculi* H3-H4 tetramer aggregation could be rescued by titration of H2A-H2B dimers. Inspection of *E. cuniculi* residue changes mapped onto the nucleosome structure suggested that this can be explained by a propensity for polymerisation via a novel H4:H4 4HB association. Such a polymerisation of histone fold heterodimers is reminiscent of the continuous *M. fervidus* histone-like helix, with the interesting implication that this behaviour could represent a reversion in *E. cuniculi* to make use of an ancestral chromatin property. This would only be possible for histones that interact weakly with DNA and each other, since the cooperativity of strong polymeric binding would otherwise render packaged DNA inaccessible.

Taken together, this work suggests that the 4 core histones with their stable and mutually exclusive 4HB H3:H3 and H4:H2B combinations generate am evolutionarily conserved histone octamer structure that can bind DNA relatively non-specifically with sufficient affinity to form a discrete nucleosome. This particle is sufficiently stable to act as a negative regulator for access to DNA binding proteins such as transcription factors yet can be displaced by polymerases and remodelling complexes. Nucleosome formation is compatible with considerable histone sequence variation which enables adaptation in DNA binding affinities to modulate chromatin stability, and changes in interaction surfaces to enable the nucleosome to act as a substrate and scaffold for genomic mechanisms. Thus, the nucleosome is a uniform building block that facilitates diversity in chromatin packaging.

## Materials and Methods

### Histone diversity calculations

Predicted proteomes of 1769 species in Ensembl Genomes release 102 (December 2020) (43) were searched by local BLAST (44) with histone fold domains (HFDs) from H2AC4 27-88, H2BC12 38-101, H3C4 64-130 and H4C1 31-92 as reference human histones based on PDB entry 2CV5 (45) and used in subsequent biochemical studies. Additional human isoforms are redundant with these reference sequences. For proteomes with multiple subspecies or isolates the proteome with highest average HFD identity was taken as representative, yielding 1208 unique species cases. Global HFD alignments were performed using EMBOSS Needle (46). Analysis was performed using python, pandas and seaborn libraries.

### Histone recombinant expression and purification

Histone genes were codon optimised for *E. coli* and synthesised (MWG Eurofins). Optimised gene sequences were flanked by NdeI and BamHI restriction sites to facilitate subcloning into pET3a expression vector. Rosetta2 pLysS cells were transformed with histone plasmids and expressed using 2YT media supplemented with IPTG. Histone inclusion body preparation and purification were carried out essentially as previously described (47).

### Histone complex refolding and purification

Purified histones were unfolded in 20 mM Tris pH 7.5, 7 M Guanidinium-HCl, 10 mM DTT at room temperature. Equimolar amounts of unfolded histones were mixed followed by dialysis against a refolding buffer 20 mM Tris pH 7.5, 2 M NaCl, 5 mM ß-mercaptoethanol at 4 ºC with 3 replenishments at 2 h intervals. *E. cuniculi* dimer:tetramer titrations were carried out at indicated ratios and NaCl concentrations by the same method. Refolded histone octamer, tetramer and dimer complexes were concentrated to 0.5 mL to 1 mL with a centrifugal concentrator then purified by Superdex 200 gel filtration chromatography followed by centrifugal concentration.

### Nucleosome assembly and native PAGE

Nucleosomes were normally assembled by mixing purified histone octamers with equimolar amounts of DNA in a solution containing 10 mM Tris-Cl pH 7.5, 2 M NaCl. Stepwise salt dialysis was carried out at 4 ºC sequentially against 0.8 M, 0.6 M, 0.5 M and 0.1 M NaCl with 10 mM Tris-Cl pH 7.5 buffer for 2 h each. Nucleosomes assembled by salt-urea dialysis used purified histones and equimolar amounts of DNA in 10 mM HEPES pH 8.0, 2 M NaCl and 1 mM EDTA dialysed against 10 mM HEPES pH 8.0, 2 M NaCl, 5 M Urea, 1 mM DTT for 12 to 16 h followed by sequentially 1.2 M, 1 M, 0.8 M, 0.6 M NaCl containing 10 mM HEPES pH 8.0, 5 M Urea and 1 mM DTT for 1 h each. Finally, the assembly was dialysed in 10 mM HEPES pH 8.0, 0.6 M NaCl, 1 mM DTT for 3 h followed by 10 mM HEPES pH 8.0, 1 mM EDTA for 3 h or overnight. Nucleosome core particles were resolved on 6% native polyacrylamide gels in 0.2 x TBE that were pre-equilibrated and run at 4ºC and stained with ethidium bromide if necessary before imaging.

### Hydroxyl radical footprinting

A 10 µL footprinting reaction sample was prepared by mixing 20-30 pmol of freshly prepared nucleosome containing Cy5 end-labelled DNA with 1 μL of 200 μM NH_4_FeSO_4_ and 400 μM EDTA mixture and incubating on ice for 15 min. Then 5 μL of 18 mM ascorbic acid in 100 mM Tris pH 7.5 was added followed by 5 μL of the 0.6 % H_2_O_2_ and incubation on ice for 90 min. After incubation, DNA was isolated by adding 20 μL of 25:24:1 phenol:chloroform:isoamyl alcohol. The aqueous phase containing DNA was transferred into a new tube supplemented with 10 μg of carrier tRNA. DNA was precipitated and purified by ethanol precipitation then resuspended in 12 μL of formamide sample loading buffer. Samples heated at 95°C for 5 min then loaded onto a hot pre-equilibrated 8 % denaturing polyacrylaminde gel with 6M urea and run at 50-55°C. Gels were imaged with a Fuji FLA5100 fluorimager.

### Thermal unfolding assay

36 pmol nucleosome or tetrasome was mixed with freshly diluted 5 x Sypro Orange in 20 µL of 20 mM Tris-Cl, pH 7.5, 100 μM NaCl in a 96 well MicroAmp Fast optical plate and thermal unfolding was observed by incubation in 1ºC increments in a StepOne Plus Thermal Cycler (24). Data was normalised for fluorescence intensity with *X. laevis* nucleosomes as a reference sample.

### Mononucleosomal DNA library preparation and SELEX

Mononucleosomal DNA was extracted from micrococcal nuclease digestion of HEK293 chromatin (gift of Dr H Dodson, NUI Galway). End repair and 5’ phosphate addition was carried out for 3 µg DNA in 1 x T4 DNA ligase buffer with 20 U T4 polynucleotide kinase, 5 U Klenow enzyme, 6 U T4 DNA polymerase and 0.4 mM dNTPs at 20 °C for 30 min. A single 3’ adenine addition was incorporated by using 1 x NEB buffer 2, 0.2 mM dATP and 15 U Klenow Exo-at 37 °C for 30 min. Ligation with adapter GTCATAGCTGTTTCCTGTGTGA with a 3’ *T overhang was carried out using T4 DNA ligase at 20 ºC for 30 min. The initial DNA library was PCR amplified in bulk using Taq polymerase and the adapter as primers for 25 cycles. Nucleosomes were assembled using a 10:1 molar excess of DNA to octamer and then separated by native PAGE as described above. Nucleosomal DNA was identified by ethidium bromide staining of native PAGE gels and by incubation of excised gel slices in 0.5 M NH_3_CH_3_COOH, 10 mM Mg(CH_3_COO)_2_, 1 mM EDTA pH 8.0 at 50 ºC for 30 min followed by centrifugation then purification by QIA-PCR clean up (Qiagen). The 193 bp DNA from each round was amplified by KOD then Taq polymerase and purified by anion exchange chromatography using a MonoQ column (48) to enable successive nucleosome assembly and selection rounds. Samples were submitted for multiplexed paired end Illumina sequencing with 100 base read length (EMBO Genomics Core). Raw reads were deposited in as NCBI BioProject accession PRJNA600132.

### SELEX data analysis

Raw reads were filtered for quality and adapters removed using Cutadapt 1.18 (49) then mapped to GRCh37.p5 using Bowtie 2.1.0 (50), yielding 145-175 bp fragments of quality ≥25. Abundances were analysed with fastaptamer-count and fastaptamer-cluster v1.0.3 (51) and poweRlaw 0.70.6 (52), and local blast (44). Periodicity was analysed using a custom implementation of the integer period discrete Fourier transform (53). Additional analysis and processing was performed using python, perl and R.

## Acknowledgments

We are grateful to Dr Helen Dodson for HEK293 mononucleosomal DNA, Dr Jonathan Doran for assistance with hydroxy radical footprinting, and the EMBO Genomics Core and Dr Vladimir Benes for sequencing. This research was funded by SFI grant 08/IN.1/B1946 to AF, a NUIG Beckman Fund award to IP and a NUIG College of Science fellowship to HT.

## Supplementary Figure Legends

Supplementary figure 1: **Histone types represented in genomes by phylogenetic grouping**. Percentage of genomes with 0-4 histones type homologues at least 30% identical to human reference sequences for 305 vertebrates, 117 non-vertebrate metazoans, 96 plants, 1014 fungi, 237 protists and all 1725 genomes.

Supplementary figure 2: **Conservation by residue across histone fold domains by histone type and phylogenetic grouping**. 21,571 histones from 1769 genomes with ≥40% identity to human reference types, indexed by human reference sequence with gaps removed and grouped as vertebrates, non-vertebrate metazoans, plants, fungi and protists for A. H2A (cyan), B. H2B (grey), C. H3 (yellow) and D. H4 (blue). Horizontal bars represent mean identity for α1 helix, L1 loop, α2 helix, L2 loop and α3 helix in each plot respectively. Consensus sequence for phylogenetic grouping below.

Supplementary figure 3: **Alignment of recombinant histones**. Histone sequences expressed in supplementary figure 4 with N-terminal methionine removed, aligned by Clustal Omega and visualised with pyBoxshade. HFD motifs as for supplementary figure 2.

Supplementary figure 4: **Purified recombinant histones:** SDS PAGE of purified recombinant histones stained with Coomassie.

Supplementary figure 5: **Formation of histone octamers by divergent histones**. A. Gel filtration chromatograms of octamer refolding with soluble aggregate (7 mL), octamer (12.5 mL) and dimer (16 mL) indicated by hatched lines. B. SDS PAGE of purified octamers from A. C. SDS PAGE of peak fractions from *L. major* refolding. H2A/H3 and H2B/H4 cannot be resolved due to similar sizes but are consistent with octamer and dimer respectively.

Supplementary figure 6: **Hydroxy radical footprinting of 601 nucleosomes assembled with divergent octamers**. A. Denaturing gel of forward strand hydroxy radical footprinting of Cy5 labelled 601 DNA assembled with recombinant octamers for *X. laevis* (Xl), human (Hs), *S. cerevisiae* (Sc), *P. falciparum* (Pf), *G. lamblia* (Gl) and *E. cuniculi* (Ec) indicating superhelical locations (SHLs) and base coordinates along the strand. Marker lane G is a G track cut at guanine nucleotides. B. Extended run of samples from A to visualise region 67-147 bp. C. Profile of indicated lanes from panels A and B.

Supplementary figure 7: **Comparison of ionic strength effect on *X. laevis* and *E. cuniculi* tetramer refolding**. A. Gel filtration chromatograms of *X. laevis* and *E. cuniculi* (H3-H4)_2_ tetramer refolded in 0.5 M, 1 M, 1.5 M and 2 M NaCl, 1 mM EDTA. Soluble aggregate (7 mL) and tetramer (14 mL) indicated by hatched lines. B. SDS PAGE of fractions from *X. laevis* chromatograms in A showing excess H3 in aggregate and equimolar H3 and H4 in tetramer. C. SDS PAGE of fractions from *E. cuniculi* chromatograms in A showing equimolar H3 and H4 in both aggregate and tetramer.

Supplementary figure 8: **Chemical nature of 4HB interface in *E. cuniculi***. A. 4HB face of human H3 (yellow) showing aliphatic hydrophobic sidechain (black), aromatic hydrophobic sidechains (grey), acidic sidechains (red) and basic sidechains (blue) with hydrophilic residue labels above and hydrophobic labels below. *E. cuniculi* differences (left, bold) are conservative in chemical nature. B. Equivalent 4HB face of human H4 in same orientation and colouring to A. Human to *E. cuniculi* differences (left, bold) shown as coloured disks involve major charge-hydrophilic (R67S, E74T; white), charge-hydrophobic (D68I; black), hydrophilic-hydrophobic (T71I; black) and hydrophobic-charge (Y88H; blue) that completely change nature of H4 interface in 4HB. C. Model for potential polymeric association of *E. cuniculi* tetramers driven by homodimerization of hydrophobic H4:H4 4HBs modulated by H2A-H2B capping.

Supplementary figure 9: **Assembly of *E. cuniculi* tetramers on MMTV-derived DNA fragments**. A. Native PAGE of *X. laevis* (lanes 2-7) and *E. cuniculi* (lanes 8-13) (H3-H4)_2_ tetramers assembled on MMTV derived 252 bp 105A0 DNA by salt-urea dialysis giving species consistent with mono-, di- and tri-tetrasomes for *X. laevis* but only mono-tetramer for *E. cuniculi* tetramer assembly. B. Titration of *E. cuniculi* (H3-H4)_2_ tetramers at indicated ratio on 147 bp MMTV nucA, nucB and nucAupstream (nucAu) DNA in lanes 4-12 showing affinity for AT-rich nucAu. Assembled *X. laevis* di-tetramers on same DNA as marker in lanes 1-3. Tetrasome assembly was carried out using salt-urea step dialysis. C. GC content of MMTV 3’LTR region showing Schematic of MMTV 3’LTR region showing DNA fragment locations from panels A and B, and GC% plot over 31 bp sliding window. 147 bp nucB, nucA and nucAu DNA fragments have overall GC% of 35%, 70% and 34% respectively.

Supplementary figure 10: **Assembly of nucleosomes on ANK1-POU6F2 fusion**. A. Native PAGE of nucleosomes assembled in 147 bp 601, nucA and ANK1-POU6F2 fusion (ANK) with human octamer, visualised by ethidium Bromide staining. B. Thermal stability assay of nucleosomes in A.

## Supplementary Table Legends

Supplementary table 1: **Properties of recombinant histones**. Histone IDs from UniProt. Molecular mass, net charge and extinction coefficient from Expasy Protpram. *H. sapiens* (Hs), *X. laevis* (Xl), *Plasmodium falciparum D7* (Pf), *Giardia lamblia ATCC 50803* (Gl), *Leishmania major Friedlin* (Lm), *Encephalitozoon cuniculi GB_M1* (Ec).

Supplementary table 2: **Properties of nucleosomes assembled from recombinant octamers on 601 DNA**. Molecular mass and net charge from Expasy Protpram. Relative mobility from figure 2.

Supplementary table 3: **Human genomic sequences selected by SELEX**. See also table 1 and supplementary table 4.

Supplementary table 4: **Properties of human genomic sequences selected by SELEX**. Abundance as counts per million total fragments (cpM) amongst human genome derived fragments at SELEX round 6. Location in human genome (GRCh37). Length range of mapped fragments for S1 and S2. Significant 10-11 bp periodic dinucleotides in fragment (p 0.05) where S is C or G. GC% and percentile rank of this GC% within SELEX input library.

## References

1. R. D. Kornberg, Chromatin structure: a repeating unit of histones and DNA. Science (New York, NY) 184, 868–871 (1974).

2. K. Luger, A. W. Mäder, R. K. Richmond, D. F. Sargent, T. J. Richmond, Crystal structure of the nucleosome core particle at 2.8 A resolution. Nature 389, 251–260 (1997).

3. D. D. Banks, L. M. Gloss, Equilibrium folding of the core histones: the H3-H4 tetramer is less stable than the H2A-H2B dimer. Biochemistry 42, 6827–6839 (2003).

4. K. Luger, T. J. Richmond, DNA binding within the nucleosome core. Current opinion in structural biology 8, 33–40 (1998).

5. A. Elbahnsi, R. Retureau, M. Baaden, B. Hartmann, C. Oguey, Holding the Nucleosome Together: A Quantitative Description of the DNA-Histone Interface in Solution. Journal of chemical theory and computation 14, 1045–1058 (2018).

6. C. A. Davey, D. F. Sargent, K. Luger, A. W. Maeder, T. J. Richmond, Solvent mediated interactions in the structure of the nucleosome core particle at 1.9 a resolution. Journal of Molecular Biology 319, 1097–1113 (2002).

7. S. Tan, C. A. Davey, Nucleosome structural studies. Current opinion in structural biology 21, 128–136 (2011).

8. B. Alberts, et al., Molecular Biology of the Cell, 6th Ed. (Garland Science, 2014).

9. J. Dai, et al., Probing nucleosome function: a highly versatile library of synthetic histone H3 and H4 mutants. Cell 134, 1066–1078 (2008).

10. S. Nakanishi, et al., A comprehensive library of histone mutants identifies nucleosomal residues required for H3K4 methylation. Nature structural & molecular biology 15, 881–888 (2008).

11. K. Matsubara, N. Sano, T. Umehara, M. Horikoshi, Global analysis of functional surfaces of core histones with comprehensive point mutants. Genes to cells : devoted to molecular & cellular mechanisms 12, 13–33 (2007).

12. D. M. Truong, J. D. Boeke, Resetting the Yeast Epigenome with Human Nucleosomes. Cell 171, 1508–1519 (2017).

13. R. D. Kornberg, Y. Lorch, Primary Role of the Nucleosome. Mol Cell 79, 371–375 (2020).

14. K. Luger, M. L. Dechassa, D. J. Tremethick, New insights into nucleosome and chromatin structure: an ordered state or a disordered affair? Nat Rev Mol Cell Biol 13, 436–447 (2012).

15. P. B. Talbert, S. Henikoff, Histone variants--ancient wrap artists of the epigenome. Nat Rev Mol Cell Biol 11, 264–275 (2010).

16. J. Bao, M. T. Bedford, Epigenetic regulation of the histone-to-protamine transition during spermiogenesis. Reproduction (Cambridge, England) 151, R55–70 (2016).

17. S. G. Gornik, et al., Loss of nucleosomal DNA condensation coincides with appearance of a novel nuclear protein in dinoflagellates. Curr Biol 22, 2303–2312 (2012).

18. F. Mattiroli, et al., Structure of histone-based chromatin in Archaea. Science (New York, NY) 357, 609–612 (2017).

19. L. Eme, A. Spang, J. Lombard, C. W. Stairs, T. J. G. Ettema, Archaea and the origin of eukaryotes. Nature Reviews Microbiology 15, 711–723 (2017).

20. L. A. Katz, Origin and diversification of eukaryotes. Annual review of microbiology 66, 411–427 (2012).

21. V. L. Koumandou, et al., Molecular paleontology and complexity in the last eukaryotic common ancestor. Critical Reviews in Biochemistry and Molecular Biology 48, 373–396 (2013).

22. C. L. White, R. K. Suto, K. Luger, Structure of the yeast nucleosome core particle reveals fundamental changes in internucleosome interactions. The EMBO journal 20, 5207–5218 (2001).

23. S. Chakravarthy, K. Luger, The histone variant macro-H2A preferentially forms “hybrid nucleosomes”. Journal of Biological Chemistry 281, 25522–25531 (2006).

24. H. Taguchi, N. Horikoshi, Y. Arimura, H. Kurumizaka, A method for evaluating nucleosome stability with a protein-binding fluorescent dye. Methods (San Diego, Calif) 70, 119–126 (2014).

25. S. Bowerman, J. Wereszczynski, K. Luger, Archaeal chromatin “slinkies” are inherently dynamic complexes with deflected DNA wrapping pathways. Elife 10, e65587 (2021).

26. K. R. Shieh, et al., AptCompare: optimized de novo motif discovery of RNA aptamers via HTS-SELEX. Bioinformatics 36, 2905–2906 (2020).

27. P. T. Lowary, J. Widom, New DNA sequence rules for high affinity binding to histone octamer and sequence-directed nucleosome positioning. 276, 19–42 (1998).

28. M. Malm, et al., Evolution from adherent to suspension: systems biology of HEK293 cell line development. Sci Reports 10, 18996 (2020).

29. A. A. Stepanenko, V. V. Dmitrenko, HEK293 in cell biology and cancer research: phenotype, karyotype, tumorigenicity, and stress-induced genome-phenotype evolution. Gene 569, 182–190 (2015).

30. E. Brookes, et al., Polycomb Associates Genome-wide with a Specific RNA Polymerase II Variant, and Regulates Metabolic Genes in ESCs. Cell Stem Cell 10, 157–170 (2012).

31. M. R. Hass, et al., Runx1 Shapes the Chromatin Landscape Via a Cascade of Direct and Indirect Targets. Biorxiv, 2020.09.25.313767 (2020).

32. R. Perales, L. Zhang, D. Bentley, Histone occupancy in vivo at the 601 nucleosome binding element is determined by transcriptional history. Molecular and cellular biology 31, 3485–3496 (2011).

33. V. Alva, A. N. Lupas, Histones predate the split between bacteria and archaea. Bioinformatics 35, 2349–2353 (2018).

34. P. B. Talbert, M. P. Meers, S. Henikoff, Old cogs, new tricks: the evolution of gene expression in a chromatin context. Nat Rev Genet 20, 283–297 (2019).

35. A. Flaus, T. J. Richmond, Positioning and stability of nucleosomes on MMTV 3’LTR sequences. 275, 427–441 (1998).

36. A. Hughes, O. J. Rando, Chromatin “programming” by sequence--is there more to the nucleosome code than %GC? Journal of Biology 8, 96 (2009).

37. D. Tillo, T. R. Hughes, G+C content dominates intrinsic nucleosome occupancy. BMC Bioinformatics 10, 442 (2009).

38. Y. Field, et al., Distinct modes of regulation by chromatin encoded through nucleosome positioning signals. PLoS Comput Biol 4, e1000216 (2008).

39. J. T. Huff, D. Zilberman, Dnmt1-independent CG methylation contributes to nucleosome positioning in diverse eukaryotes. Cell 156, 1286–1297 (2014).

40. E. Silberhorn, et al., Plasmodium falciparum Nucleosomes Exhibit Reduced Stability and Lost Sequence Dependent Nucleosome Positioning. PLoS pathogens 12, e1006080 (2016).

41. P. R. Kensche, et al., The nucleosome landscape of Plasmodium falciparum reveals chromatin architecture and dynamics of regulatory sequences. Nucleic acids research 44, 2110–2124 (2015).

42. T. Hollin, M. Gupta, T. Lenz, K. G. L. Roch, Dynamic Chromatin Structure and Epigenetics Control the Fate of Malaria Parasites. Trends Genet 37, 73–85 (2020).

43. A. D. Yates, et al., Ensembl 2020. Nucleic Acids Res 48, D682–D688 (2019).

44. C. Camacho, et al., BLAST+: architecture and applications. Bmc Bioinformatics 10, 421 (2009).

45. Y. Tsunaka, N. Kajimura, S. Tate, K. Morikawa, Alteration of the nucleosomal DNA path in the crystal structure of a human nucleosome core particle. Nucleic acids research 33, 3424– 3434 (2005).

46. P. Rice, I. Longden, A. Bleasby, EMBOSS: The European Molecular Biology Open Software Suite. Trends Genet 16, 276–277 (2000).

47. K. Luger, T. J. Rechsteiner, T. J. Richmond, Preparation of nucleosome core particle from recombinant histones. Methods Enzymol 304, 3–19 (1999).

48. M. Bruno, A. Flaus, T. A. Owen-Hughes, Site-specific attachment of reporter compounds to recombinant histones. Methods Enzymol 375, 211–228 (2004).

49. M. Martin, Cutadapt removes adapter sequences from high-throughput sequencing reads. Embnet J 17, 10–12 (2011).

50. B. Langmead, S. L. Salzberg, Fast gapped-read alignment with Bowtie 2. Nat Methods 9, 357–359 (2012).

51. K. K. Alam, J. L. Chang, D. H. Burke, FASTAptamer: A Bioinformatic Toolkit for High-throughput Sequence Analysis of Combinatorial Selections. Mol Ther - Nucleic Acids 4, e230 (2015).

52. C. S. Gillespie, Fitting Heavy Tailed Distributions: The poweRlaw Package. J Stat Softw 64, 1–16 (2015).

53. J. Epps, H. Ying, G. A. Huttley, Statistical methods for detecting periodic fragments in DNA sequence data. Biology direct 6, 21 (2011).

