## Supplementary Figures and Tables for "Histone sequence variation in divergent eukaryotes facilitates diversity in chromatin packaging"

Supplementary Figure 1

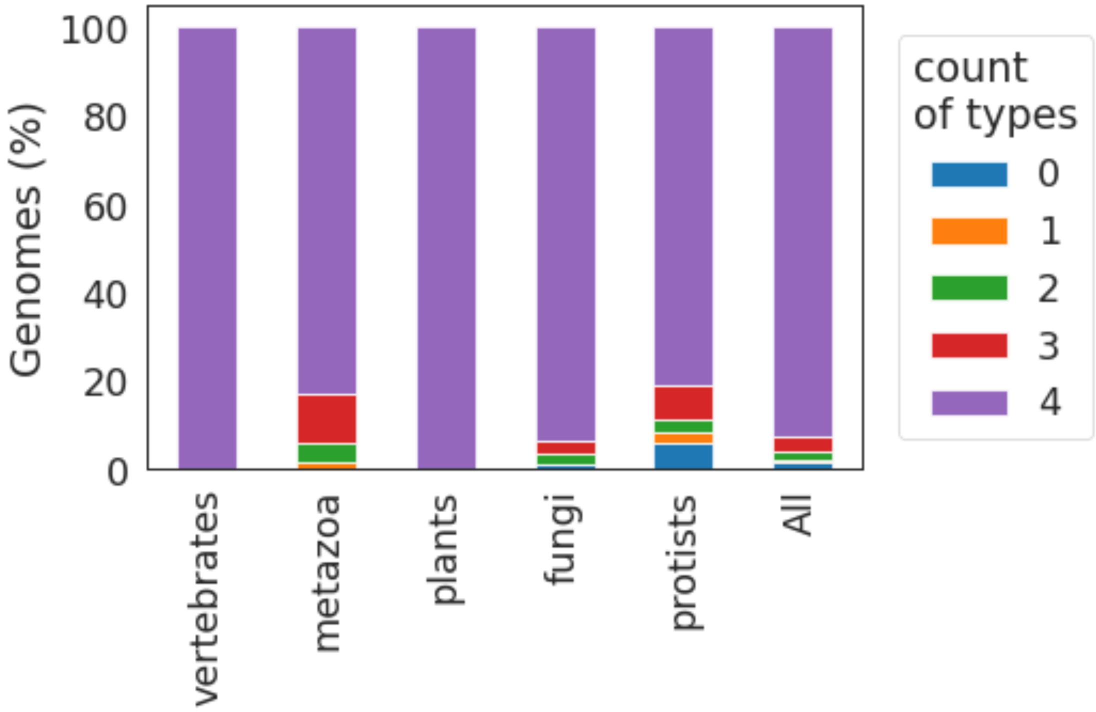

Supplementary Figure 2

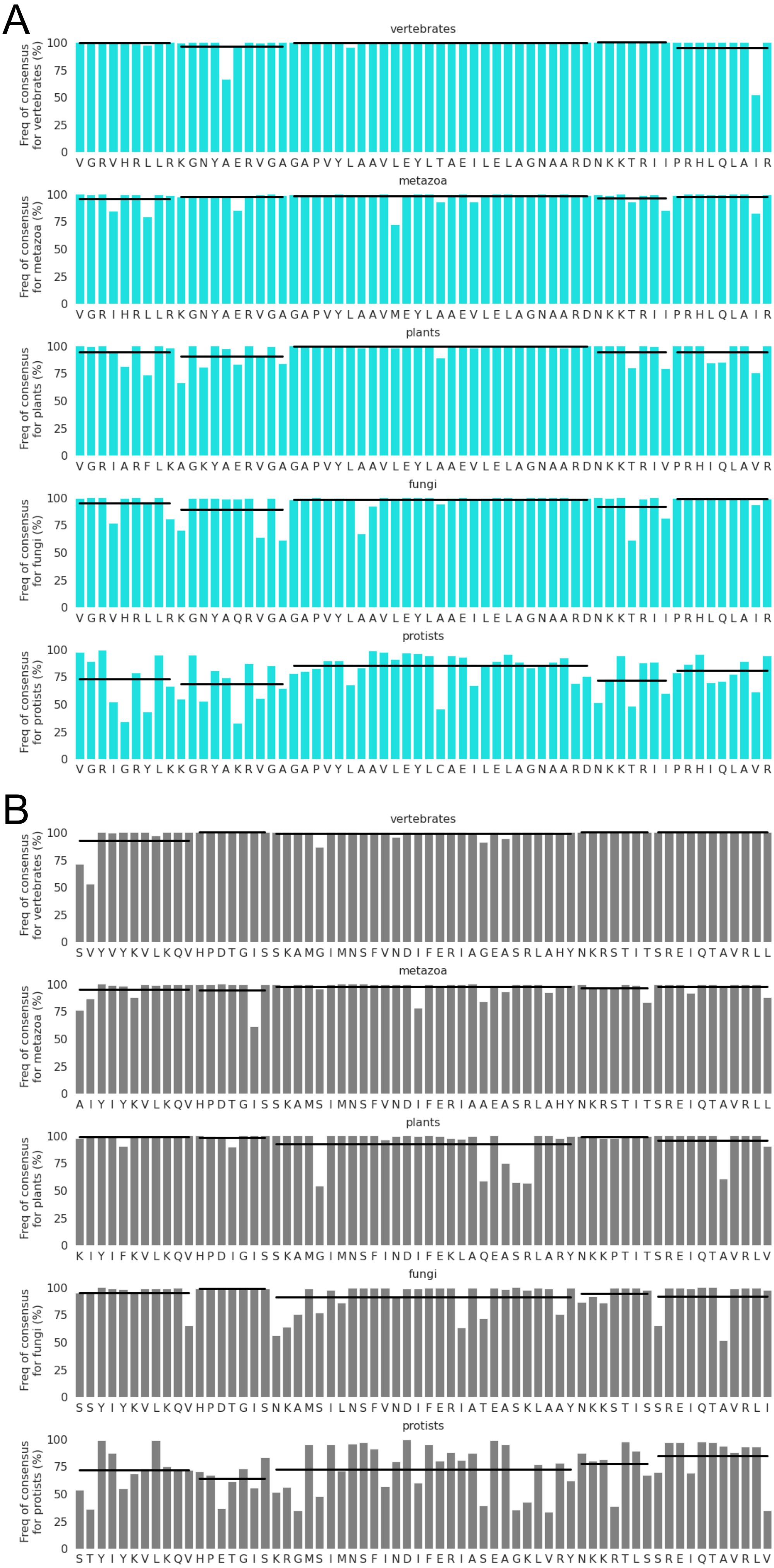

C

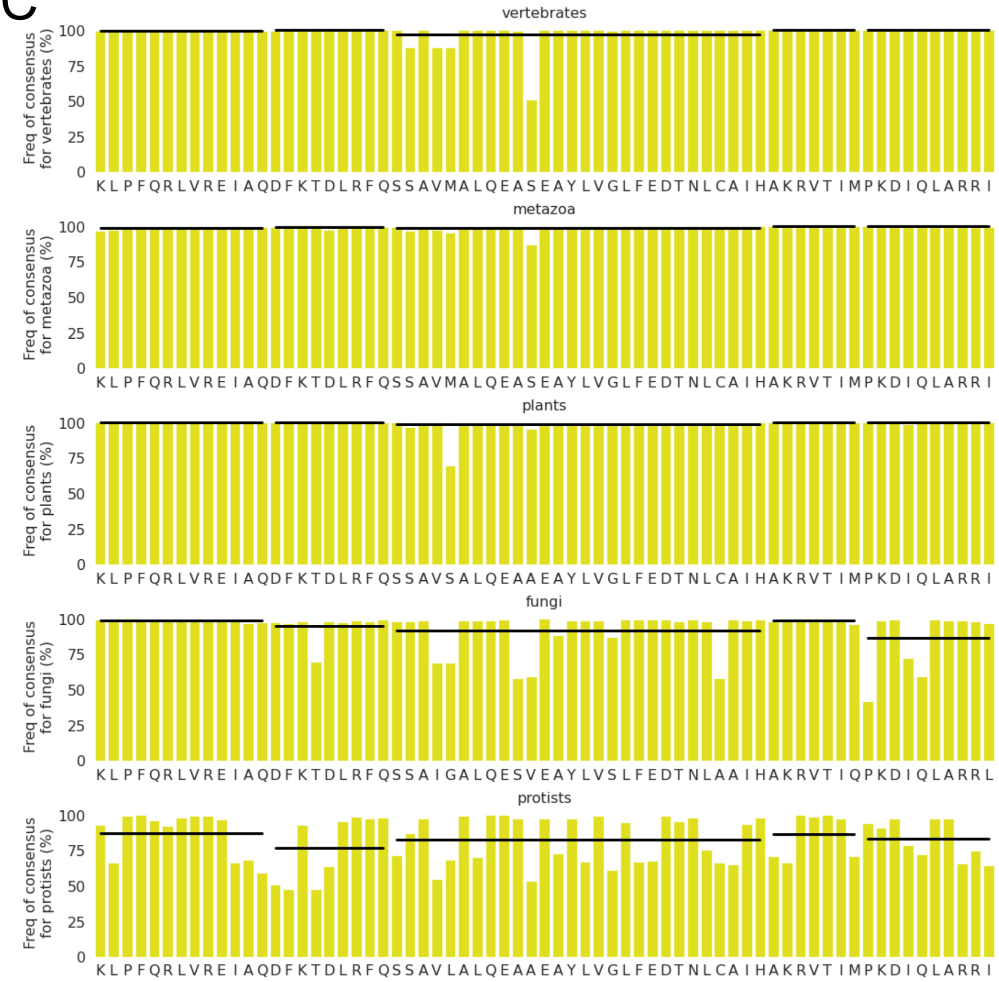
vertebrates
Freq of consensus for vertebrates (%)
0 25 50 75 100
KLPFQRLVREIAQDFKTDLRFQSSAVMALQEASEAYLVGLFEDTNLCAIHAKRVTIMPKDIQLARRI
metazoa
Freq of consensus for metazoa (%)
0 25 50 75 100
KLPFQRLVREIAQDFKTDLRFQSSAVMALQEASEAYLVGLFEDTNLCAIHAKRVTIMPKDIQLARRI
plants
Freq of consensus for plants (%)
0 25 50 75 100
KLPFQRLVREIAQDFKTDLRFQSSAVSALQEAAEAYLVGLFEDTNLCAIHAKRVTIMPKDIQLARRI
fungi
Freq of consensus for fungi (%)
0 25 50 75 100
KLPFQRLVREIAQDFKTDLRFQSSAIGALQESVEAYLVSLFEDTNLAAIHAKRVTIQPKDIQLARRL
protists
Freq of consensus for protists (%)
0 25 50 75 100
KLPFQRLVREIAQDFKTDLRFQSSAVLALQEAAEAYLVGLFEDTNLCAIHAKRVTIMPKDIQLARRI

D

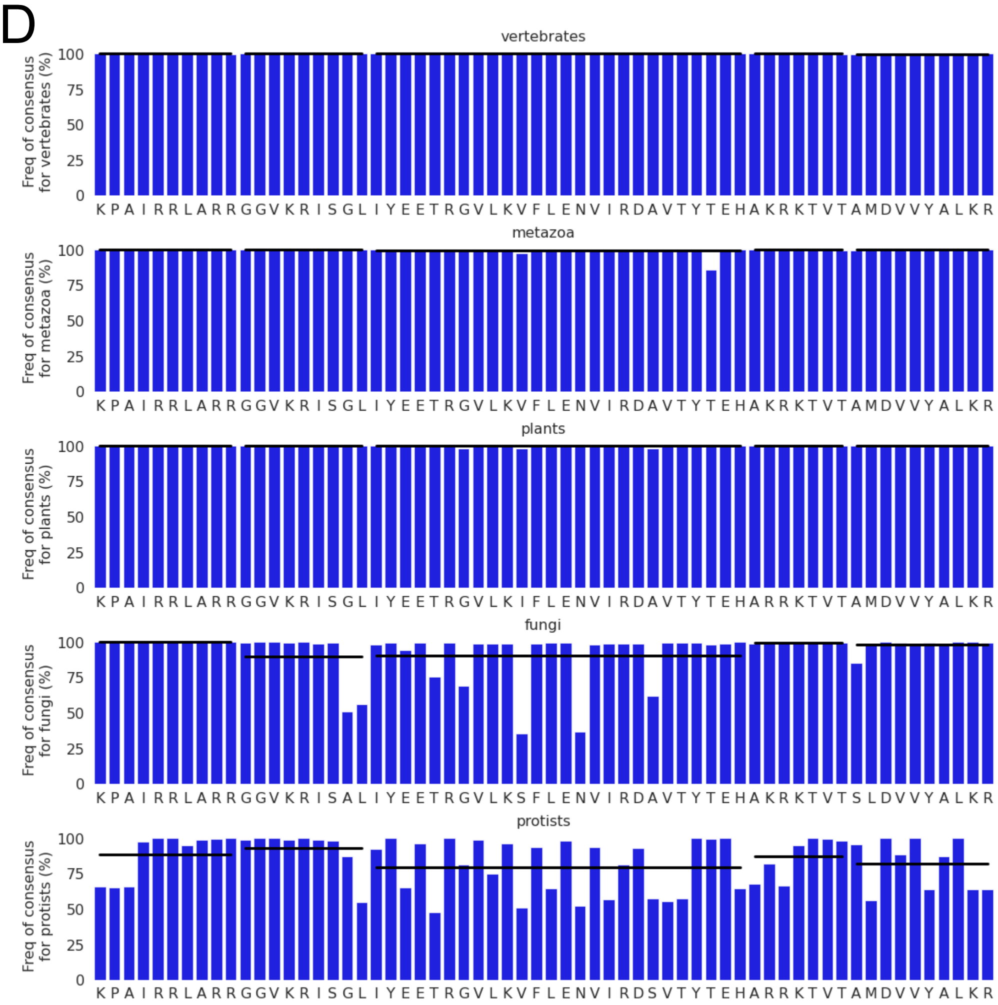
vertebrates
Freq of consensus for vertebrates (%)
0 25 50 75 100
KPAIRRLARRGGVKRISGLIYEETRGVLKVFLENVIRDAVTYTEHAKRKTVTAMDVVYALKR
metazoa
Freq of consensus for metazoa (%)
0 25 50 75 100
KPAIRRLARRGGVKRISGLIYEETRGVLKVFLENVIRDAVTYTEHAKRKTVTAMDVVYALKR
plants
Freq of consensus for plants (%)
0 25 50 75 100
KPAIRRLARRGGVKRISGLIYEETRGVLKIFLENVIRDAVTYTEHARRKTVTAMDVVYALKR
fungi
Freq of consensus for fungi (%)
0 25 50 75 100
KPAIRRLARRGGVKRISALIYEETRGVLKSFLENVIRDAVTYTEHAKRKTVTSLDVVYALKR
protists
Freq of consensus for protists (%)
0 25 50 75 100
KPAIRRLARRGGVKRISGLIYEETRGVLKVFLENVIRDSVTYTEHARRKTVTAMDVVYALKR

Supplementary Figure 3

A

```
HsH2A ---SGRGKQGGKARAKAKTRSSRAGLQFPVGRVHRLLRKGNYSERVGAGAPVYLAAVLEYLTAEILELAGNAAR-
XlH2A ---SGRGKQGGKTRAKAKTRSSRAGLQFPVGRVHRLLRKGNYAERVGAGAPVYLAAVLEYLTAEILELAGNAAR-
ScH2A --SGGKGGKAGSAAKASQSRSAKAGLTFPVGRVHRLLRRGNYAQRIGSGAPVYLTAVLEYLAAEILELAGNAAR-
PfH2A ---SAKGKTGRKKASKGTSNSAKAGLQFPVGRIGRYLKKGKYAKRVGAGAPVYLAAVLEYLCAEILELAGNAAR-
LmH2A ----ATPRSAKKAARKSSTKSAKAGLIFPVGRVGGMMRRGQYARRIGASGAVYLAAVLEYLTAELLELSVKAAAQ
GlH2A ------STKPVKDNSKMKSRSARAGISFPIGRIHRHLREGRYAERISSDAPVYLAAVLENVVAEVFREACNHRD-
EcH2A VVIQGKGGKADP--RVIGKDEEHQKSIVKLSQIKKI-MKDRTRMRISKDALVAVSACVMYLISEITDGAKNVAS-
                                        α1          L1                α2

HsH2A -DNKKTRIIPRHLQLAIRNDEELNKLLGRVTIAQGGVLPNIQAVLLPKKTESHHKAKGK--
XlH2A -DNKKTRIIPRHLQLAVRNDEELNKLLGRVTIAQGGVLPNIQSVLLPKKTESSKSAKSK--
ScH2A -DNKKTRIIPRHLQLAIRNDDELNKLLGNVTIAQGGVLPNIHQNLLPKKSAKTAKASQE-L
PfH2A -DNKKSRITPRHIQLAVRNDEELNKFLAGVTFASGGVLPNIHNVLLPKKSQLKAGTANQDY
LmH2A SGKKRCRLSPRTVMLAARHDNDIGTLLKSVTLSHSGVVPNISKAMAKKKGGKKGKATPSA-
GlH2A -KKSQKRIVPNHILTALRKDKELATIFANVTIREGGVARSAKEGREGKGS-----HRSQDL
EcH2A -TDGKKKVMPKHINNAICNDTELHFVGHDWLIKNGGMKSYIAPGDFAVSSKK-GSSR--D-
         L2     α3
```

B

```
HsH2B -----PEPAKSAPAPKKGSKKAVTKAQKKDGKKRKRSRKESYSVYVYKVLKQV----------HPDTGISSKAMG
XlH2B --------AKSAPAPKKGSKKAVTKTQKKDGKKRRKTRKESYAIYVYKVLKQV----------HPDTGISSKAMS
ScH2B SSAAEKKPASKAPAEKKP--AAKKTSTSVDGKKRSKVRKETYSSYIYKVLKQT----------HPDTGISQKSMS
PfH2B -------------VSKKPAKAKKTGTGPDGKKKRKKSRYDSYGLYIFKVLKQV----------HPDTGISRKSMN
LmH2B ----------------------ASSRKASNPHKSHRKPKRSWNVYVGRSLKAI----------NAQMSMSHRTMK
GlH2B -----------SKVE-TKRLMKKTEAGDKGDAKRKHKRHETYATYIYKVLRSEN----IRSEADTDLGISNKGME
EcH2B -------------AKSARHVTGKAPSSLDAHDRKKSKKKSSSMCVGSMFKSAVKRISREVSPDNNIMLTSNSIQ
                                                 α1                    L1   α2

HsH2B IMNSFVNDIFERIAGEASRLAHYNKRSTITSREIQTAVRLLLPGELAKHAVSEGTKAVTKYTSAK-----
XlH2B IMNSFVNDVFERIAGEASRLAHYNKRSTITSREIQTAVRLLLPGELAKHAVSEGTKAVTKYTSAK-----
ScH2B ILNSFVNDIFERIATEASKLAAYNKKSTISAREIQTAVRLILPGELAKHAVSEGTRAVTKYSSSTQA---
PfH2B IMNSFLVDTFEKIATEASRLCKYTRRDTLSSREIQTAIRLVLPGELAKHAVSEGTKAVTKFTSSK------
LmH2B IVNSYVNDVMERICTEAASIVRANKKRTLGAREVQTAVRIVLPAELAKHAMAEGTKAVSSASR-------
GlH2B VMNSLVNDLFERIASEASNLAKISKRNTIGKKDIESAAKLVIPGEIGRLIRDEADKALSKFTSSKESTKK
EcH2B VLCGISYDFVDTVADLAGELARKVGKQTVGSDDIISAFEILLKGELRKLAIREVHNALSKSQAKSRGK--
              α2             L2     α3
```

C

```
HsH3 ARTKQTARKST-GGKAPRKQLATKAARKSAPATGGVK-KPHRYRPGTVALREIRRYQKSTELLIRKLPFQRLVRE
XlH3 ARTKQTARKST-GGKAPRKQLATKAARKSAPATGGVK-KPHRYRPGTVALREIRRYQKSTELLIRKLPFQRLVRE
ScH3 ARTKQTARKST-GGKAPRKQLASKAARKSAPSTGGVK-KPHRYKPGTVALREIRRFQKSTELLIRKLPFQRLVRE
PfH3 ARTKQTARKST-AGKAPRKQLASKAARKSAPISAGIK-KPHRYRPGTVALREIRRYQKSTDLLIRKLPFQRLVRE
LmH3 SRTKETAR--------AKRTITSKKSKKAPSAVSGVKMSHRRWRPGTCAIREIRKFQKSTSLLIQCAPFQRLVRE
GlH3 ARTKHTARKTTSATKAPRKTIARKAARKTASSTSGIK-KTGRKKQGMVAVKEIKKYQKSTDLLIRKLPFSKLVRD
EcH3 ARTKQSARKTT-GGKAPRKQLSAKSARKGVSPASSAGAKKSRYRPGSVALKEIRRYQKSTDFLIRRLPFQRACRS
                                                                          α1

HsH3 IAQ--DFKTDLRFQSSAVMALQEACEAYLVGLFEDTNLCAIHAKRVTIMPKDIQLARRIRGERA-------
XlH3 IAQ--DFKTDLRFQSSAVMALQEASEAYLVALFEDTNLCAIHAKRVTIMPKDIQLARRIRGERA-------
ScH3 IAQ--DFKTDLRFQSSAIGALQESVEAYLVSLFEDTNLAAIHAKRVTIQKKDIKLARRLRGERS-------
PfH3 IAQ--DYKTDLRFQSSAVMALQEAAEAYLVGLFEDTNLCAIHAKRVTIMPKDIQLARRIRGERS-------
LmH3 VSS--AQKEGLRFQSSAIMALQEATEAYIVSLMADTNLACIHAKRVTIQPKDIQLALRLRGERH-------
GlH3 IVTSGLSKSDIRFQGAAVEALQESAENYIISLFVDTQLCAEHAKRVTIMKPDMELATRIGKRIEPEYRKGK
EcH3 VVKECSNATDIRFQGPALASIQEALEVYLVGLFEDAMLCAYHAKRVTVFPKDISLVLKLRSRHVKSISD--
          L1              α2             L2     α3
```

D

```
HsH4 ---SGRGKGGKGLGKGGAKRHRKVLRDNIQGITKPAIRRLARRGGVKRISGLIYEETRGVLKVFLENVIRDAVTY
XlH4 ---SGRGKGGKGLGKGGAKRHRKVLRDNIQGITKPAIRRLARRGGVKRISGLIYEETRGVLKVFLENVIRDAVTY
ScH4 ---SGRGKGGKGLGKGGAKRHRKILRDNIQGITKPAIRRLARRGGVKRISGLIYEEVRAVLKSFLESVIRDSVTY
PfH4 ---SGRGKGGKGLGKGGAKRHRKILRDNIQGITKPAIRRLARRGGVKRISGLIYEEIRGVLKVFLENVIKDSIMY
GlH4 ---SGK---GKGKGYGKSKRHS-KEKDTLGGITKPAIRRLARRGGVKRISSTIYQQTREVLKAFLEVVLRDSLTY
LmH4 ----AKGK-RSTDAKGSQRRQKKVLRDNIRGITRGSVRRMARRGGVKRISSEVYEEVRRVLKAYVEDIVRCSTAY
EcH4 NTQSIGAKGKSKAAKGIAKRHRK-QSSLSDSISKPAIRRIARRAGVRRVGGCFKEINNAAREYIRDTLSIACIY
                                         α1       L1              α2

HsH4 TEHAKRKTVTAMDVVYALKRQGRTLYGFGG
XlH4 TEHAKRKTVTAMDVVYALKRQGRTLYGFGG
ScH4 TEHAKRKTVTSLDVVYALKRQGRTLYGFGG
PfH4 TEHAKRKTVTAMDIVYSLKRQGRTLYGFGG
GlH4 TEHGQRKTVTSQDVVYALKRQGRTLYGFGI
LmH4 TEYARKKTVTACDVVNALRKQGHILYGYA-
EcH4 ATHAKRKTITCSDILHSLKRMGIKYIGY--
        L2       α3
```

Supplementary Figure 4

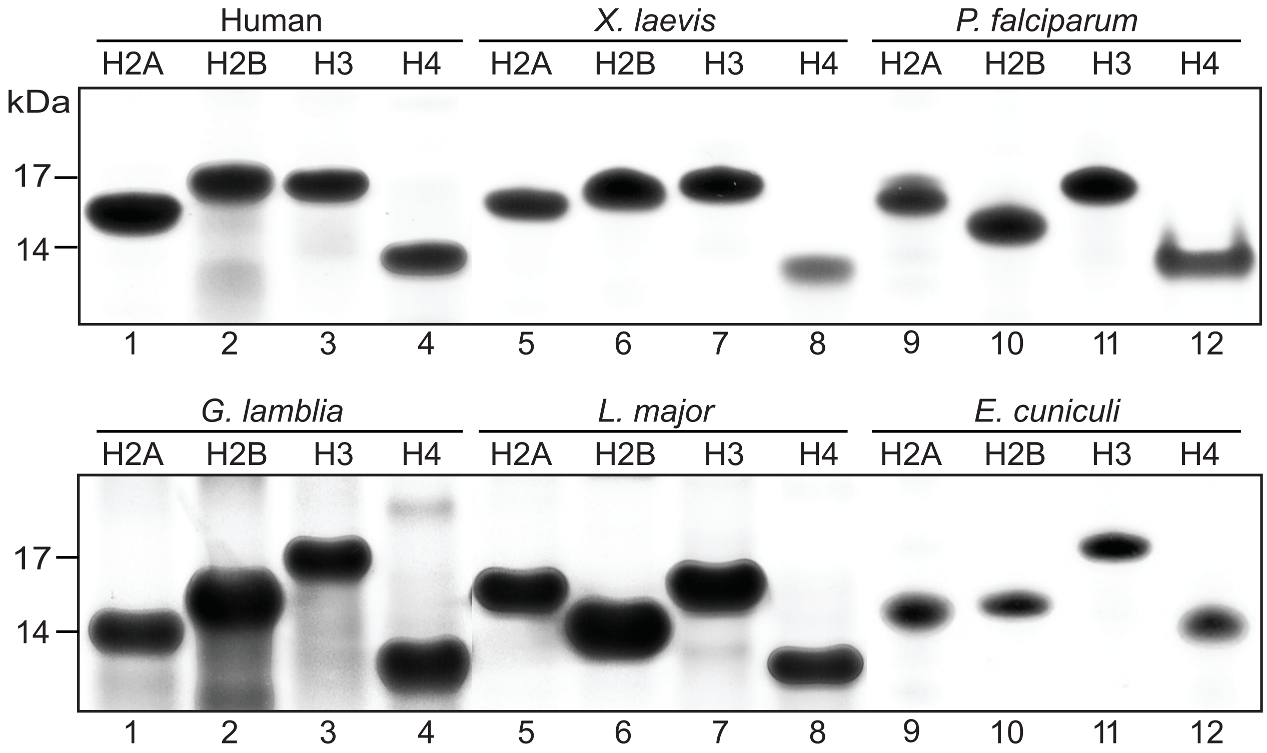

Supplementary Figure 5

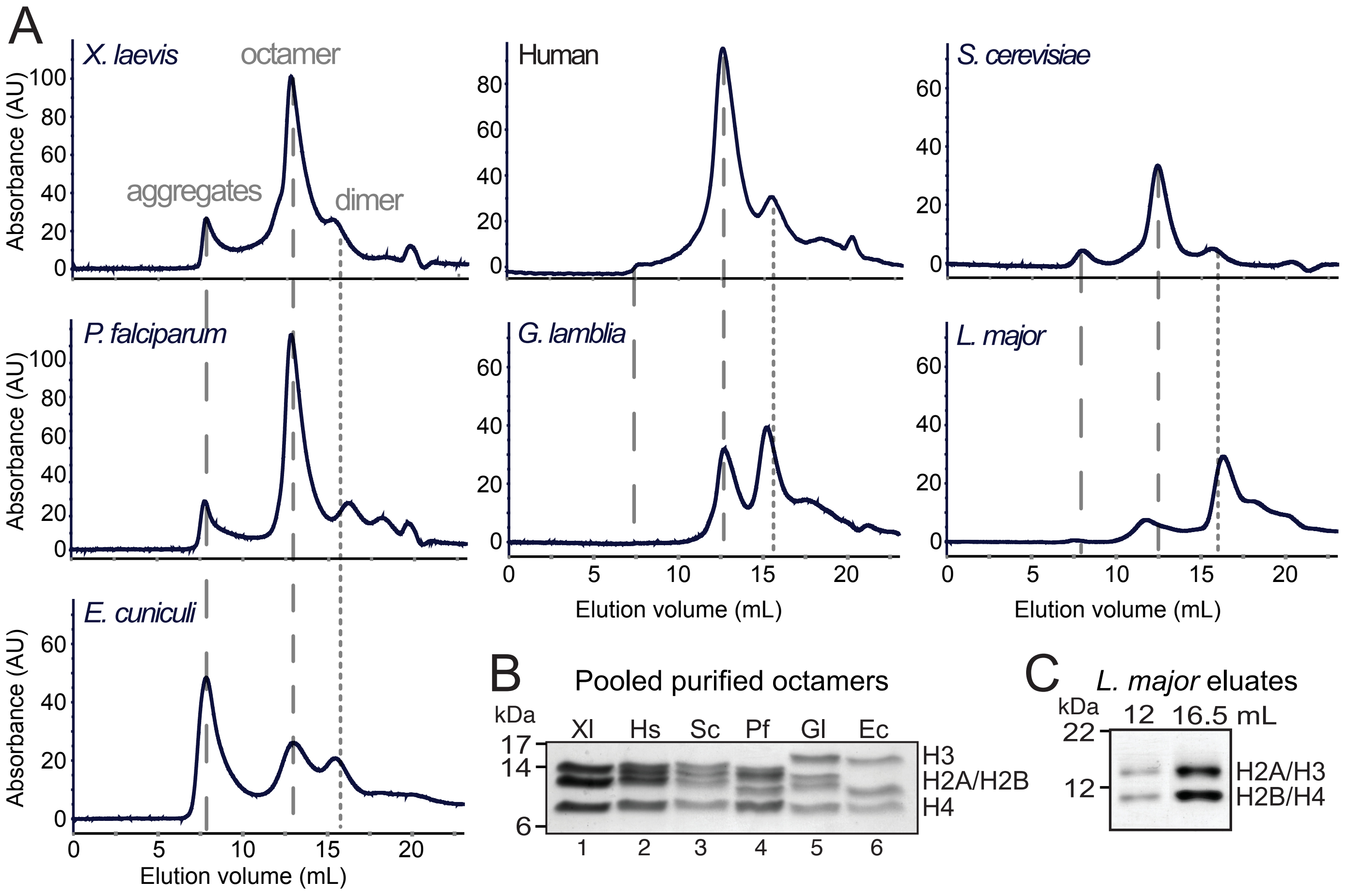

Supplementary Figure 6

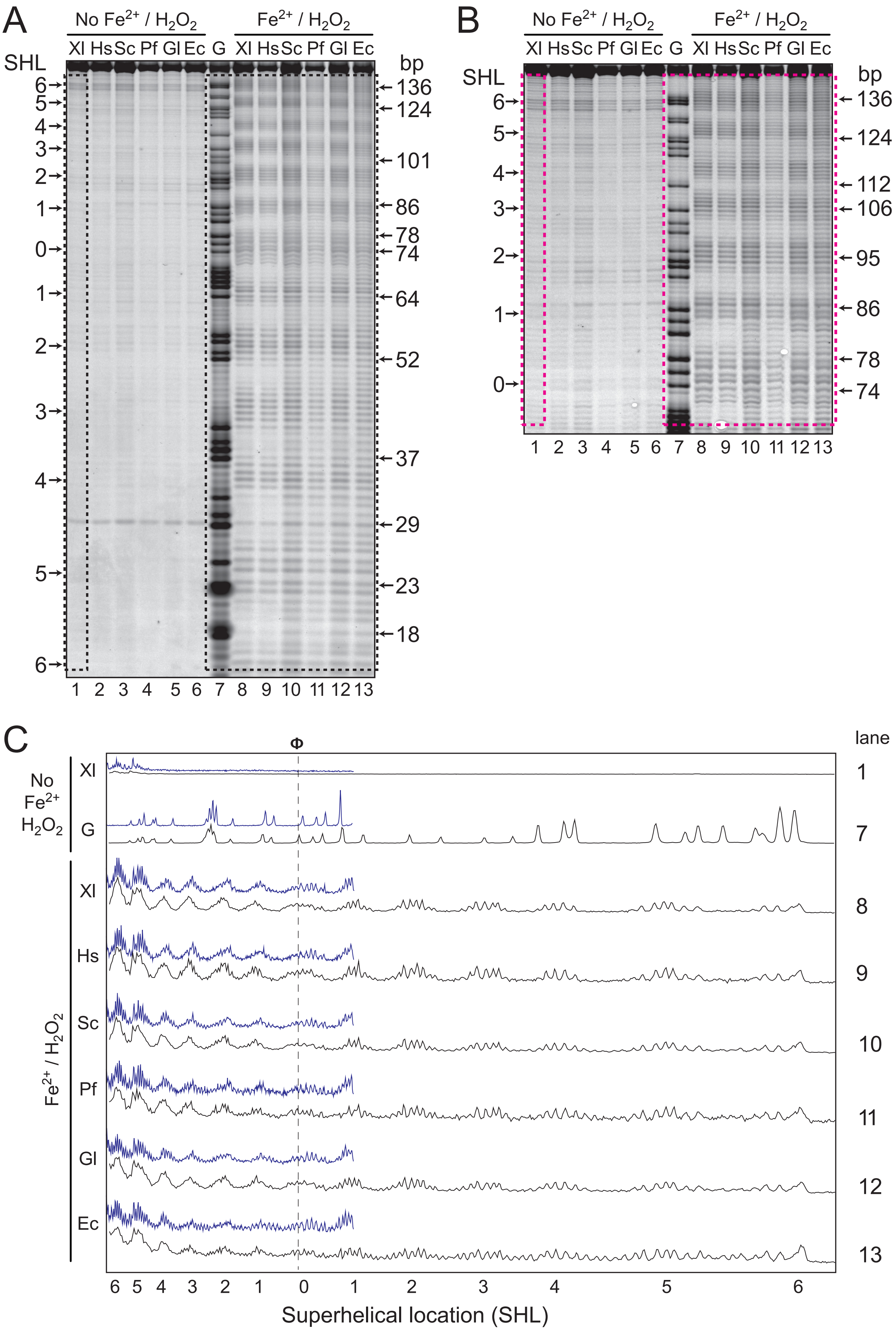

Supplementary Figure 7

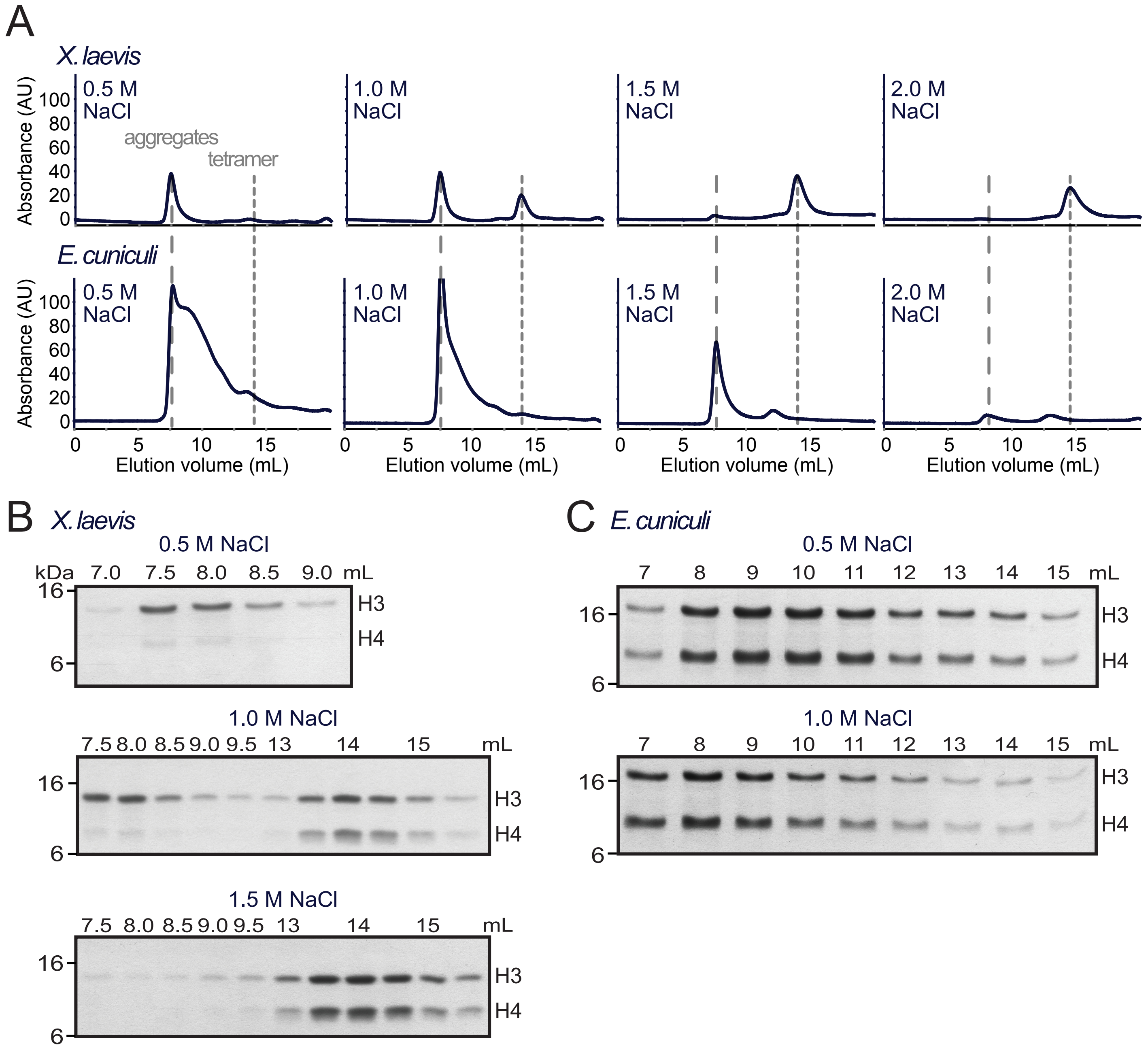

Supplementary Figure 8

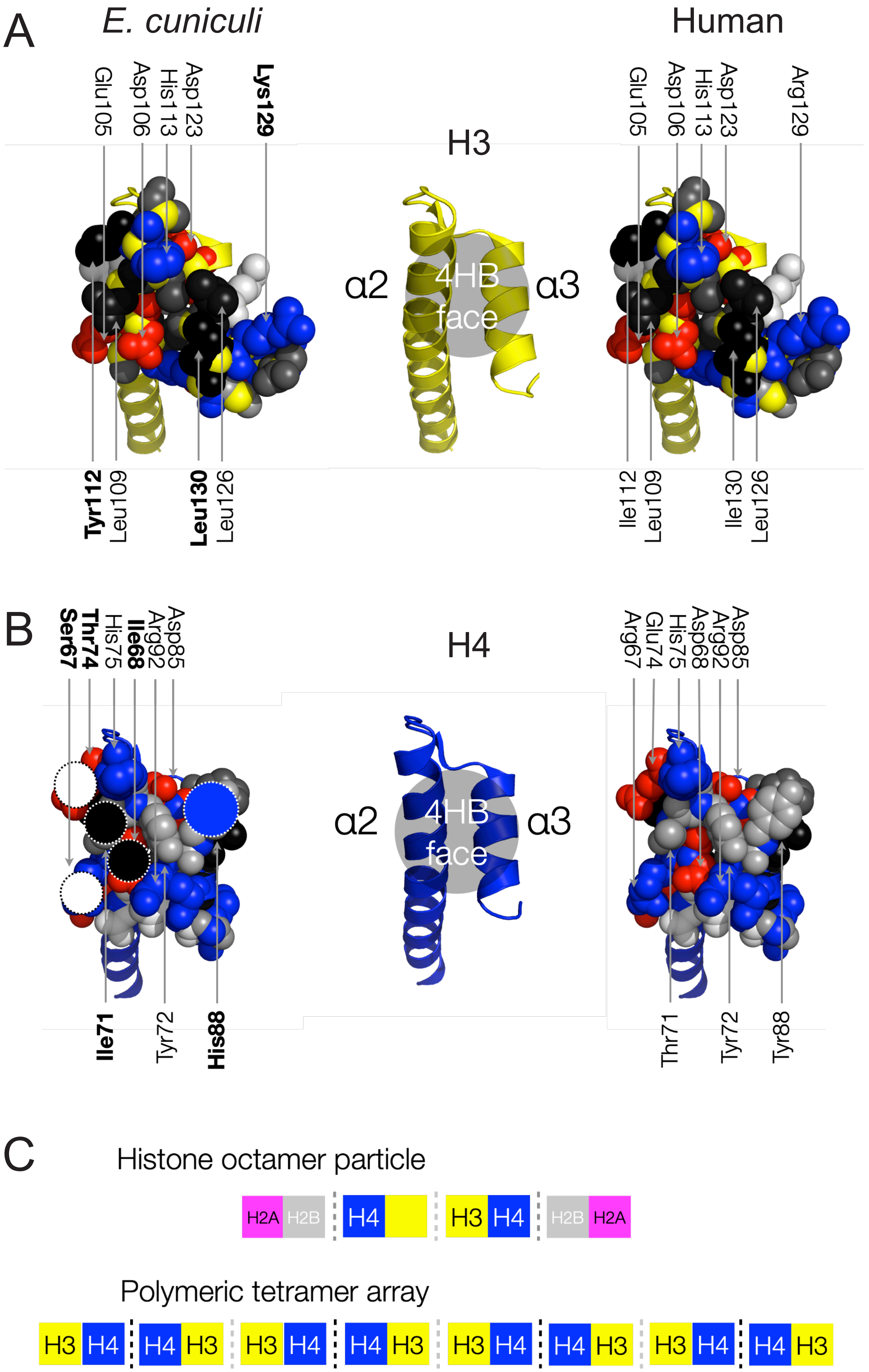

Supplementary Figure 9

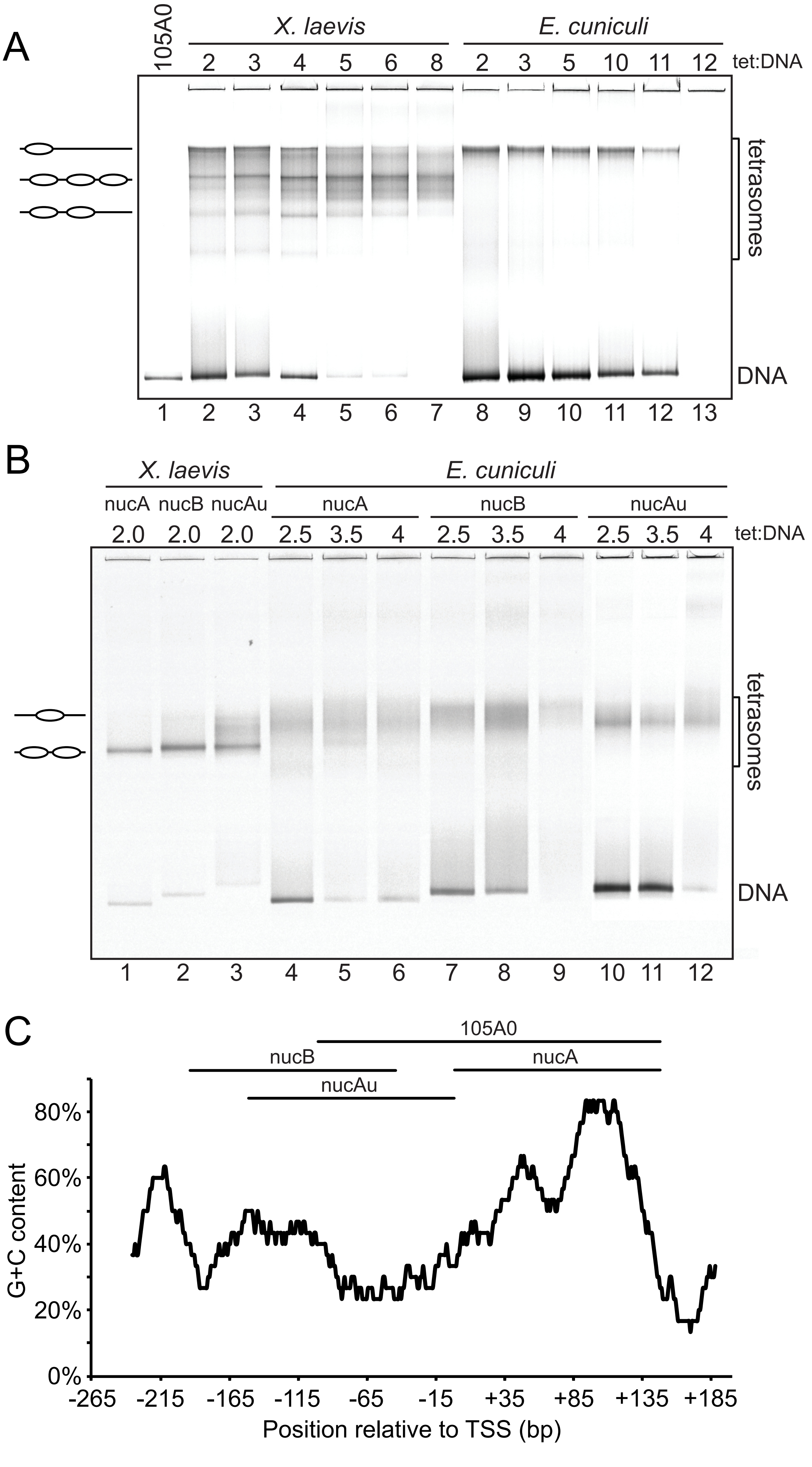

Supplementary Figure 10

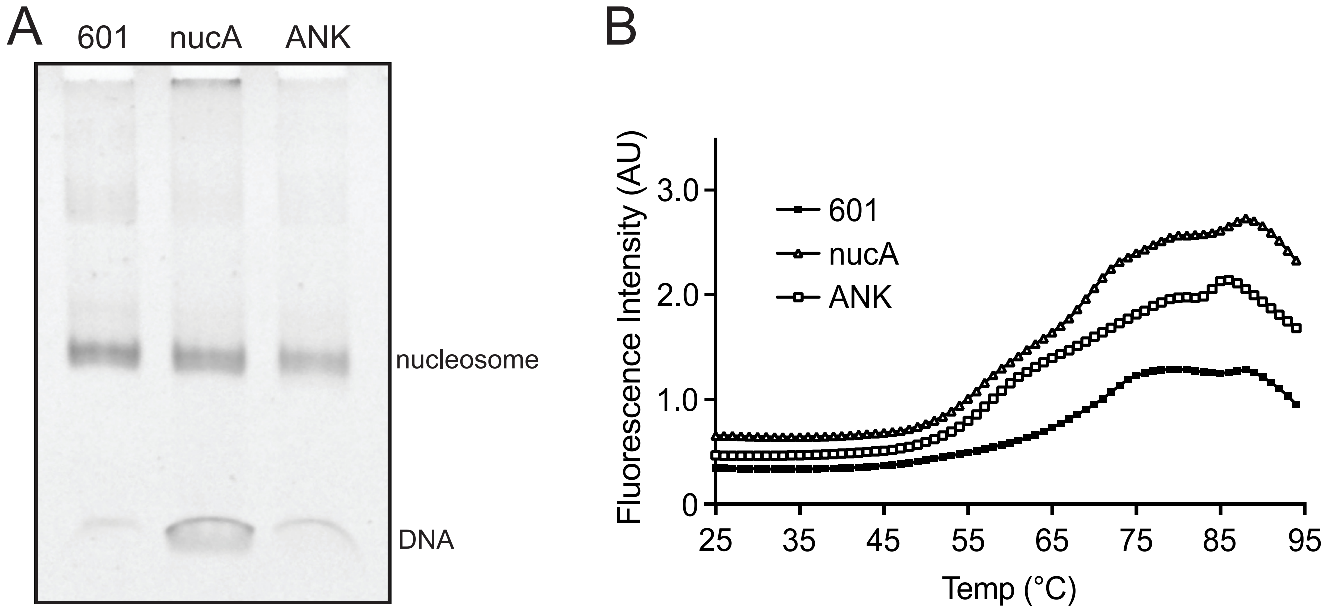

Supplementary table 1: **Properties of recombinant histones.** Histone IDs from UniProt. Molecular mass, net charge and extinction coefficient from Expasy Protpram. *H. sapiens* (Hs), *X. laevis* (Xl), *Plasmodium falciparum D7* (Pf), *Giardia lamblia ATCC 50803* (Gl), *Leishmania major Friedlin* (Lm), *Encephalitozoon cuniculi GB_M1* (Ec).

| | Histone ID | Mass (kDa) | Net charge | Extinction coefficient ($M^{-1}$ $cm^{-1}$) |
|---|---|---|---|---|
| HsH2A | H2AC4 | 14.14 | +17.9 | 4470 |
| HsH2B | H2BC12 | 13.89 | +18.6 | 7450 |
| HsH3 | H3C4 | 15.40 | +20.3 | 4470 |
| HsH4 | H4C1 | 11.37 | +18.4 | 5960 |
| XlH2A | H2AC14.L | 14.08 | +17.4 | 4470 |
| XlH2B | H2BC11.S | 13.63 | +19.6 | 7450 |
| XlH3 | H3C14.L | 15.40 | +20.4 | 4470 |
| XlH4 | H4C1.L | 11.37 | +18.4 | 5960 |
| PfH2A | PF3D7_0617800 | 14.12 | +16.4 | 7450 |
| PfH2B | PF3D7_1105100 | 13.13 | +17.4 | 5960 |
| PfH3 | PF3D7_0610400 | 15.45 | +20.4 | 5960 |
| PfH4 | PF3D7_1105000 | 11.46 | +18.4 | 5960 |
| GlH2A | GL50803_0027521 | 13.88 | +13.1 | 2980 |
| GlH2B | GL50803_00121045 | 14.59 | +9.4 | 4470 |
| GlH3 | GL50803_00135231 | 16.32 | +22.4 | 4470 |
| GlH4 | GL50803_00135002 | 11.10 | +16.4 | 7450 |
| LmH2A | LMJF_29_1730 | 13.82 | +22.4 | 4470 |
| LmH2B | LMJF_19_0030 | 11.85 | +15.8 | 8480 |
| LmH3 | LMJF_10_0870 | 14.68 | +18.5 | 6990 |
| LmH4 | LMJF_15_0010 | 11.44 | +17.1 | 8940 |
| EcH2A | ECU02_0720 | 14.01 | +9.8 | 8480 |
| EcH2B | ECU08_0410 | 14.06 | +11.6 | 1490 |
| EcH3 | ECU03_1460 | 15.99 | +20.3 | 5960 |
| EcH4 | ECU09_0440 | 11.38 | +19.5 | 5960 |

Supplementary table 2: **Properties of nucleosomes assembled from recombinant octamers on 601 DNA.** Molecular mass and net charge from Expasy Protpram. Relative mobility from figure 2.

| Octamer in nucleosome | Octamer mass (kDa) | Octamer net charge | Relative mobility in native PAGE (Rf) |
|---|---|---|---|
| *X. laevis* | 108.8 | +152 | 3.60 |
| *H. sapiens* | 109.5 | +151 | 3.48 |
| *S. cerevisiae* | 109.8 | +138 | 3.50 |
| *P. falciparum* | 108.2 | +146 | 3.65 |
| *G. lamblia* | 111.6 | +123 | 3.15 |
| *L. major* | 103.4 | +148 | - |
| *E. cuniculi* | 110.8 | +125 | 3.19 |

Supplementary table 3: **Human genomic sequences selected by SELEX**. See also table 1 and supplementary table 4.

| ID | Sequence |
|---|---|
| S1 | GCGGGGTCTGAACCCCGGCACGGTGAACTCCTGCTGCATTCCCACCAAGCTGAGCACCATGTCCATGCTGTACTTCGATGATGAGTACAACATCGTCAAGCGGGAC GTGCCCAACATGATTGTGGAGGAGTGCGGCTGCGCCTGACAGTGCAAGGCAGGGGCACGGTGGT |
| S2 | TACACATTCACATGTGTAAACACACAGGTGGTGACTCATACACACATGCATGCATGTGCACACAGCACCAGGGGGCCTGGCTGGCCTGTAGTGGGTGGGATGTGTG CGTTTGCCCATCAGCGACAGTGTCTCCGCCTGTCTTTGGGTGTGATCCTGGG |
| S3 | GCTGATGTGTGCTGTGTGCGCTGTGTGCTGTGTGCTGATGTGTGCTGATGTGTGCTGTGTGCTGACGTGTGCTGTGTGTGCGCTGATGTGTGCTGTGTGTGTGCTG TGTGCTGTGTGTGCGCTGATGTGTGCTGTGTGTGTGCTGTGTGCTG |
| S4 | ACCAGCTGCACCTGTCTGTAGGAGCACGCTGGTAGCATACGGAGCAGCTGCTTGCTCTCCTGCTGCCTGCAGATTCCCCGGCTGTGTCTTGTTACTTCCCTGCAAC TCCTCAGTGCCCAGGCAGATGTCTGGCTCGTGATCGGTTCCCG |
| S5 | TGCAGTCCTGCAATCACCTCCCAAATAAACCATGTGCCCTTGCTGGAATACTTGTCTCTGGGTCTGCGTCTGGGCACCTGCACGGCACGTGCTTGTGTGTTTCTCT CAGCTGATTTTGTTCCCACATGGGCAGAGTCAAGCTTGGTTTCCT |
| S6 | CCGCAGGATCCACAGCACGGAGCAAACAGCATTTGGAAGTCGTGTTCGCAGTACTTCCGGCCTTCAAACTGCAAAGGGGTCGCAGAGAGAGGACAAGCAAACCACC TATCATCCCCATGAGCAGCCCAGGCCCTTGTTGGGTCAGGACCTGACAGCAGGGGATGAGGCA |
| S7 | AGCAGCAGCAGCGTGGTGACTCCTCGTTCCTGAGCTGCTGTAGTGAGAGGGCAGACTCCGCGAGCAGCGGCTGCCTGGTCCTCTTGGCTCTGCACCGCTTTCTCCC CCTGTGAGTTCTGCCTTCTGTCGGTTTCAGCGTCTCTACCACCACAGCTTGTTGTGATCCTCCGG |
| S8 | CCTCATGGGTCACACTCTGAACTTCACCTCTAGGGACCTGCTGGCACTTGAGTCTAGTTACTGCCATAGAAGCAAAGAGTGGGTGTTGGGGCTGGGTCCTGAGGAG AAGCAGCATTGTCTTGCCTGGCAACTGCCTCAGGGGGACCATCACTGTTTCCT |
| S9 | GGCAGCGCCATCACCATGTCCTTAAACCATCCTTACAGTCACTACCCTGCATCCTTTACCCCTTTTCCCCATTCACTGTGCTCACCCAGAGAAACTCCAAACTCCA AACTCCAGTTGGATCCAACTATGGCTCAAGAACCAGCACCCCTGTGTGACT |
| S10 | GGTGTTTTGGCCACCATCACCCTCAGCCACTTCTGACAGCTGGAGCCCTACCCTGAATCTCGTTACGTGAAATCCCTCACTCATGCACTCAATGAGACATGTTGGG AATTGTGCTGGGATGTCCATCATCCTCTCCCATTAGCTGTTCCCTGCA |
| S11 | AGGGAACAGCTATCTCTCTGCAACATGGGAATTTCTCTGCAACTTGCCCTTTTTGCTCTGCACTGAGAGGTCCCTCCGTGAGGGGCGTCTGTGACTCCAGCTCCAC CTGCACTGCCACGTGACGCTCCCAGGCCGAGGAGACAACGCTGTGCATGATGGCCTCAGGGCC |
| S12 | TGTGTGCTGTGTGCCCCGTGTGTGCACATGTGCGCCCCGTGTGTGCTGTGTGACCCGTGTGCTTGTGTGTACCGTGTGTGAGCCATGTGCTGCGTGTGCTGTATGC TGTGTGCTCCGTGTGTTCTGTGCACTGTGTGTGCCCTGCG |
| S13 | AGGAACAGCTCCAGTCTGCAACGCCCAGCAAGATCAACGCAGTAGGCAGGTGATTGCTGCATTTTCAACTAGGTACACGGCTCATATCGTTGGGACTGGTTAGACA GTGGGTGCAGCCCATGGAGGGTGAGCTGAAGCAGGGTGAGGCATTGCC |
| S14 | GCGTGCGCGTGCGTGTGCCGTGCGTGTGCCGTGTGGGTGCCGTGTGCGCGTGCGTGTGCCATGTGGATGCCATGTGCCTTGCATGTGCCGTGTGGATGCCGCATGC CCGTGCGTGTGCCATGTGGATGCCGTGTGCCCGTGTGTCCACG |
| S15 | TGACATGAAATGGCAATGAGCACAGGGCCTGGTGCACAAGAATGGCCCATGTGAAGTGTGAGGGGTTGTTCACGTTCATCTGCTGTCACTTCTATCCTAGCCTAAA ACTGTTAGGAGGGGAAGGATGAGTGTGGTGTGATGCCCAAGGC |
| *ANK1-POU6F2* | AGCACAGGTGCCACACAAGACTTCAATAAAGCTTTCAAGTATCTAAGCCCACCTGGAAGTTATTGACGAACCCATGCAGATTACTGCGCAGGTCCACTCTCCACAA TTTTTATGTGTGATCCGCAGG |

Supplementary table 4: **Properties of human genomic sequences selected by SELEX.** Abundance as counts per million total fragments (cpM) amongst human genome derived fragments at SELEX round 6. Location in human genome (GRCh37). Length range of mapped fragments for S1 and S2. Significant 10-11 bp periodic dinucleotides in fragment ($p \leq 0.05$) where S is C or G. GC% and percentile rank of this GC% within SELEX input library.

| ID | Human reads (cpM) | *P. falciparum* reads (cpM) | *E. cuniculi* reads (cpM) | Chromosomal location | Associated gene | Sequence location in gene | Length (bp) | Periodic dinucleotides | GC% | Input GC% percentile rank |
|---|---|---|---|---|---|---|---|---|---|---|
| S1 | 904,796 | 999,613 | 767,233 | 2 | *INHBB* | Exon | 162-170 | TC | 60.0 | 89 |
| S2 | 93,072 | 173 | 229,581 | 5 | *PSD2* | Intron | 158-173 | - | 55.7 | 70 |
| S3 | 104 | 39 | 21 | 10 | *CFAP46* | Intron | 153 | GC,CT | 56.2 | 73 |
| S4 | 16 | 3 | 764 | 13 | *PLUT* | Exon | 149 | CC,GG,SG | 59.7 | 88 |
| S5 | 5 | 2 | 235 | 14 | *LINC00523* | Exon | 152 | - | 52.6 | 49 |
| S6 | 194 | 0 | 17 | 2 | *LIMS2* | Exon | 150 | AT/TA | 57.4 | 78 |
| S7 | 5 | 0 | 7 | 7 | - | - | 171 | - | 60.8 | 91 |
| S8 | 4 | 0 | 13 | 10 | - | - | 159 | - | 54.7 | 64 |
| S9 | 3 | 0 | 382 | 17 | *TMEM92* | Intron | 156 | - | 51.0 | 34 |
| S10 | 0 | 0 | 94 | 1 | - | - | 154 | - | 51.9 | 43 |
| S11 | 0 | 0 | 50 | 22 | - | - | 169 | - | 59.2 | 86 |
| S12 | 0 | 0 | 11 | 17 | *RBFOX3* | Intron | 146 | - | 60.3 | 90 |
| S13 | 0 | 0 | 8 | 14 | *GPHN* | Intron | 154 | - | 55.2 | 67 |
| S14 | 0 | 0 | 7 | 10 | - | - | 149 | CC,GG,CS,GC/CG | 67.8 | 99 |
| S15 | 0 | 0 | 6 | 5 | - | - | 149 | - | 51.0 | 35 |
| *ANK1-POU6F2* | 36,346 | 1,436 | 224,747 | 8,9 | *ANK1,POU6F2* | Intron, Intron | 128 | nd | 46.5 | 7 |
